# Rapid 40S scanning and its regulation by mRNA structure during eukaryotic translation initiation

**DOI:** 10.1101/2021.12.08.471614

**Authors:** Jinfan Wang, Carlos Alvarado, Byung-Sik Shin, Jonathan Bohlen, Thomas E. Dever, Joseph D. Puglisi

## Abstract

How the eukaryotic 43S preinitiation complex scans along the 5′ untranslated region (5′UTR) of a capped mRNA to locate the correct start codon remains elusive. Here, we directly track yeast 43S-mRNA binding, scanning, and 60S subunit joining by real-time single-molecule fluorescence spectroscopy. Once engaged with the mRNA, 43S scanning occurs at >100 nucleotides per second, independent of multiple cycles of ATP-hydrolysis by RNA helicases. The scanning ribosomes can proceed through RNA secondary structures, but 5′UTR hairpin sequences near start codons drive scanning ribosomes at start codons back in the 5′ direction, requiring rescanning to arrive once more at a start codon. Direct observation of scanning ribosomes provides a mechanistic framework for translational regulation by 5′UTR structures and upstream near-cognate start codons.

**One Sentence Summary:** Direct observation of scanning eukaryotic ribosomes establishes a quantitative framework of scanning and its regulation.

## Main Text

Translation initiation is tightly regulated to define the identity and quantity of the protein products encoded by 7-methylguanosine (m^7^G)-capped messenger RNAs (mRNAs) (*1*–*5*). During initiation, the 40S ribosomal subunit forms a 43S pre-initiation complex (PIC) with eIF1, eIF1A, eIF3, eIF5 and the eIF2•GTP•Met-tRNA_i_ ternary complex. The 43S PIC is recruited to the mRNA and must locate the correct start codon, forming a 48S PIC with the start codon base paired to the anticodon of Met-tRNA_i_ in the ribosomal peptidyl site (P site). This is followed by the joining of the catalytic 60S subunit to assemble an 80S complex competent for polypeptide elongation. Extensive biochemical, structural and genetic analyses suggest that the 43S PIC binds near the 5′end of a capped mRNA facilitated by a multi-protein eIF4F complex and moves directionally through the 5′ untranslated region (5′UTR) in search of the first encountered start codon, in a process called scanning (**Fig. 1A**) (*1, 6, 7*). Initiation in cells occurs in seconds to minutes (*8*–*10*), and scanning ribosomes must navigate a range of 5′UTR lengths with secondary structures that generally impair translation (*11*–*14*). Translation is stimulated by RNA helicases (e.g. eIF4A and Ded1p in yeast) that couple putative RNA unwinding activities to ATP hydrolysis (*15*–*20*), but their precise action during scanning and initiation remains enigmatic. Here we apply direct real-time single-molecule analysis of eukaryotic translation initiation, monitoring the speed, directionality, and energy requirements of ribosomal scanning, and how scanning is modulated by 5′UTR length and structures. Finally, we explore how scanning is coupled to start-site recognition and its interplay with mRNA structures.

**Fig. 1.**
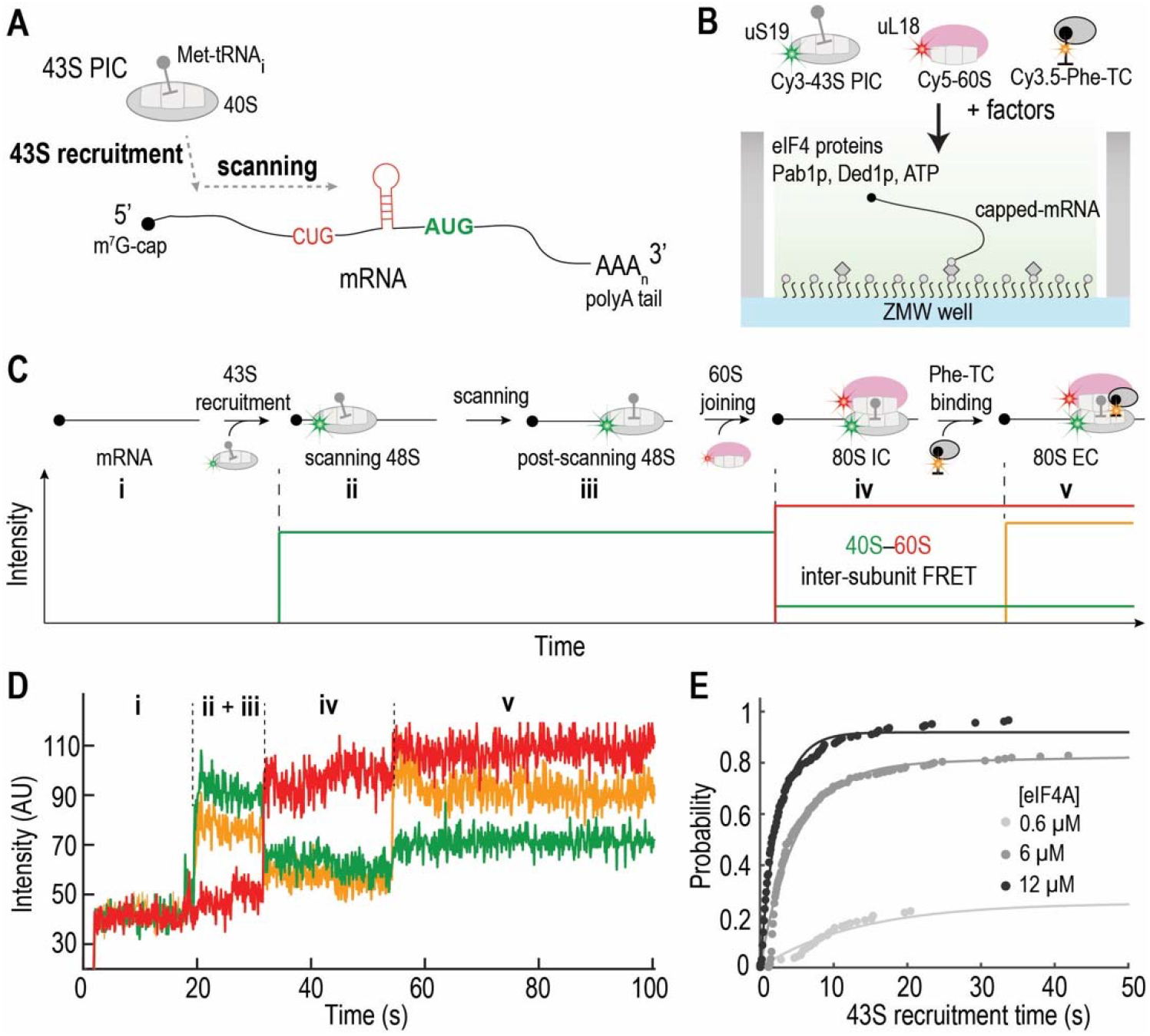
Real-time detection of initiation on single mRNAs. (**A**) Schematic (protein factors omitted) of the molecular events in early initiation. (**B**) Single-molecule assay setup. Capped-mRNA was tethered in zero-mode waveguides (ZMWs) and preincubated with a set of factors, and fluorescently labeled ribosomes and elongator TC were delivered along with factors to the ZMWs to start the reaction (see **fig. S1A**, Scheme 2). Schematic (**C**) and example trace (**D**) showing 43S PIC (Cy3, green; direct illumination) binding to unlabeled mRNA, 60S joining (Cy5, red; FRET with Cy3), and the acceptance of the first elongator tRNA (Cy3.5, yellow; direct illumination) by the 80S. (**E**) Cumulative probability distributions of the observed 43S recruitment times at different eIF4A concentrations. Data were fit to a single- or double-exponential equation to estimate the mean 43S recruitment times (data are shown in **fig. S2, C and E)**.

### Observing translation initiation in real time

We established a real-time *in vitro* single-molecule fluorescence assay (*21*) to monitor yeast 43S recruitment and initiation on surface-immobilized capped mRNAs (**Fig. 1B, fig. S1 and S2**). Upon 532-nm laser excitation, reactions were started with the addition of fluorescently labeled ribosomal subunits along with other requisite factors to *RPL30* mRNAs, and movies were collected at 30ºC and 10 frames per second (fps). 43S–mRNA binding was detected as the appearance of Cy3-40S signal (Cy3 at the N-terminus of uS19). Subsequent joining of Cy5-60S (Cy5 at the C-terminus of uL18) led to Cy3–Cy5 inter-subunit Förster resonance energy transfer (FRET) with Cy3-40S. Transition to elongation by the 80S was signified by the fluorescence of the first elongator Cy3.5-Phe-tRNA (**Fig. 1, C and D**). Controls experiments demonstrated that these signals were indeed tracking canonical cap-dependent initiation (**fig. S2A**).

Pre-incubating immobilized *RPL30* mRNAs with eIF4 proteins (eIFs 4A, 4B, 4E and 4G), Pab1p and Ded1p greatly hastened 43S recruitment (**fig. S2, B, C and E**). Rapid 43S recruitment (< 10 s at 40 nM 43S) was driven by high concentrations of eIF4A (near-saturating at 6 to 12 μM) and its ATP hydrolysis, but not Ded1p (**Fig. 1E, fig. S2, C-E**). Changes in concentrations of eIF4A (0.6 to 12 μM) or 60S (100 to 200 nM) had minor effects on 60S joining (∼30 s), while omitting Ded1p slowed the 60S joining by nearly twofold (∼56 s) (**fig. S2D and E**). These concentration changes did not affect the transition from initiation to elongation (**fig. S2B and E**), consistent with eIF5B governing this step (*21, 22*). Collectively, the RNA helicases eIF4A and Ded1p plausibly play different roles in regulating the rate-limiting steps between 43S–mRNA binding and 60S joining, i.e. scanning, and/or rearrangements post scanning (**Fig. 1C**).

To dissect the early steps of initiation and to observe scanning directly, we established a FRET signal (**Fig. 2A and B, fig. S3**) between Cy5-labeled C-terminus of uS5 (at the 40S mRNA entry channel) and a Cy3-dye located in the 3′ region of the AUG (position +20, with A of the AUG as 0). The distance between these sites should decrease during scanning and the FRET efficiency (*E*_FRET_) therefore reports on the relative 40S location on the mRNA. Combined with the uS19–uL18 inter-ribosomal subunit FRET, these two orthogonal FRET signals from a single ribosome would differentiate between the scanning and post-scanning states (**Fig. 2A**).

**Fig. 2.**
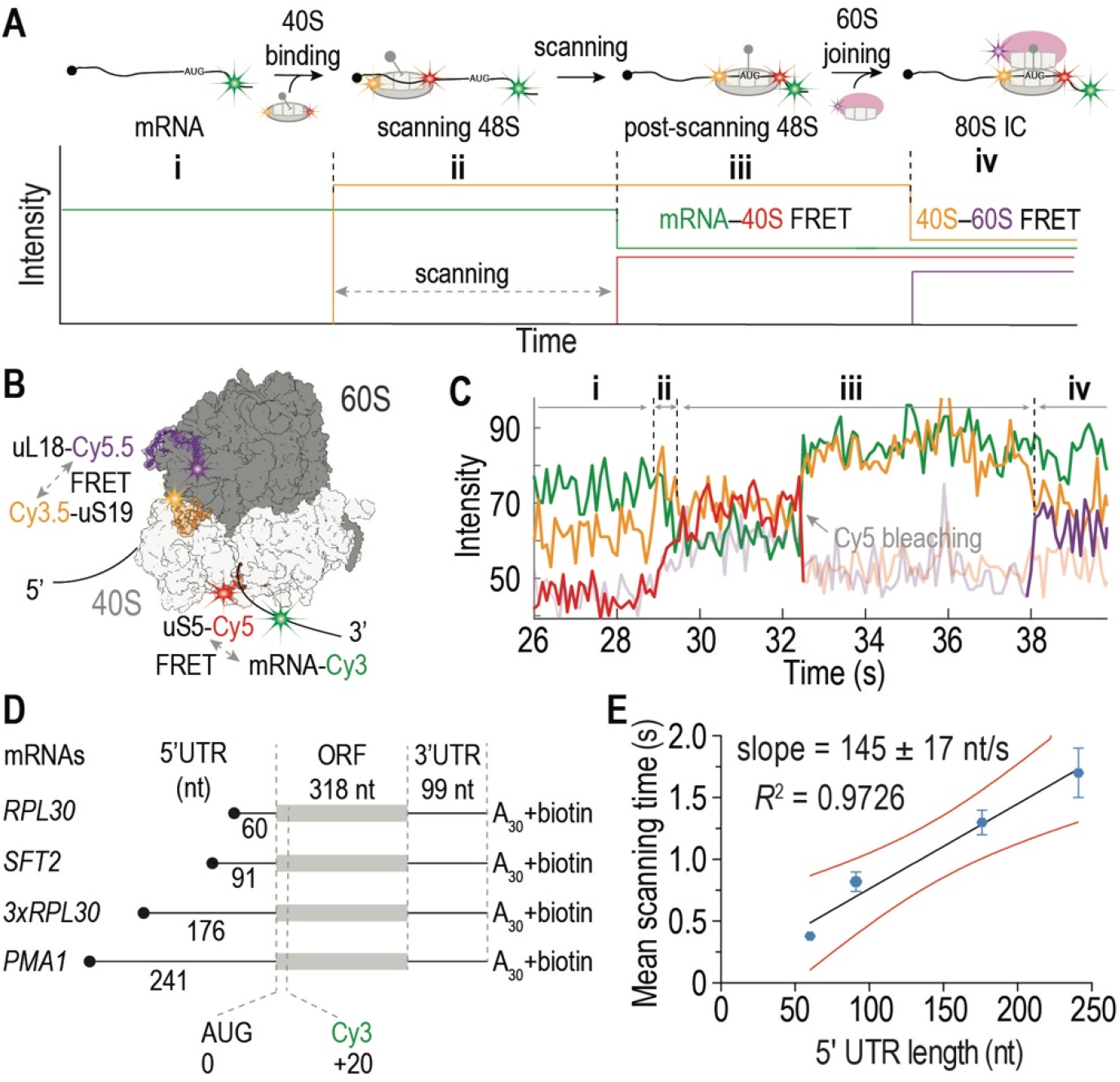
Probing scanning using two orthogonal FRET pairs with four-color detection. (**A**) Schematics of the experimental design to probe scanning by two-orthogonal FRET pairs. The 43S recruitment is detected by direct illumination of the Cy3.5 dye (yellow) on uS19, scanning is monitored by the FRET between the Cy5 dye (red) on uS5 and the Cy3 dye (green) on the mRNA downstream of the start codon, and 60S joining will result in the inter-subunit FRET between Cy3.5-uS19 and uL18-Cy5.5 (purple). (**B**) Structural model showing the labeling sites on the ribosomal subunits and mRNA. (**C**) Example trace showing 40S (Cy3.5, yellow) binding to mRNA (Cy3, green), a delay to the appearance of the mRNA–uS5 (Cy5, red) FRET, and the subsequent 60S (Cy5.5, purple) joining by Cy3.5–Cy5.5 FRET (see also **fig. S4A**). (**D**) The assayed mRNAs of varying 5′UTRs. (**E**) The linear relationship between the mean scanning time and the 5′UTR length. Data are shown in **fig. S4C**. Errors and red curves represent 95% confidence intervals (CI).

Using a similar reaction scheme and condition as above (**Fig. 1B**), we observed rapid 43S recruitment as expected (mean time ∼3.8 s), typified by the burst of the Cy3.5-uS19 signal from direct excitation by a 532-nm laser (**Fig. 2C** and **fig. S4**). After 43S recruitment, there was a brief lag (∼0.3 s, mean scanning time) until the Cy3-Cy5 *E*_FRET_ reached its maximum. This was followed by another dwell time (∼29 s, mean post-scanning state lifetime) before 60S joining, signaled by the appearance of the Cy3.5-uS19 to Cy5.5-uL18 inter-subunit FRET. Thus, this new experimental setup resulted in a similar timescale of 60S joining after 43S recruitment (∼ 0.3 + 29 s) to that determined above (∼30 s) and suggested that scanning on *RPL30* was rapid (∼0.3 s).

### Scanning is fast and its timescale is dependent on the 5′UTR length

To determine the 5′UTR length-dependence of scanning time, we tested a wide range of yeast 5′UTR lengths (*23*). We mutated the *RPL30* mRNA to contain three tandem repeats of the native *RPL30* 5′UTR, or the native 5′UTRs of *SFT2* or *PMA1* genes (**Fig. 2D**). The mean scanning time increased linearly with the 5′UTR length, corresponding to a scanning rate of 145 ± 17 nt•s^-1^ (**Fig. 2E**). By contrast, mean post-scanning state lifetime and 5′UTR length were not correlated (**fig. S4D**).

The linear relationship between scanning time and the number of nucleotides scanned by the ribosome was also observed within the *PMA1* mRNA when the Cy3-dye was installed at varying positions within the 5′UTR using Cy3-DNA oligonucleotides (corresponding to a scanning rate of 141 ± 12 nt•s^-1^, **fig. S5, A to D**). Interestingly, the anti-sense DNA oligonucleotide was not dislodged during scanning (**fig. S5, E and F**), and instead led to stalling of the scanning ribosomes (**fig. S5G**). These experiments also suggested that the 43S landing site was close to the 5′end of the mRNA, as mRNA-uS5 FRET was observed with the donor dye on the mRNA positioned proximal to the 5′ end while distal to the start site.

### Scanning is globally 5′ to 3′ unidirectional

To obtain more robust mechanistic details about the scanning process, we examined FRET solely between uS5-Cy5-40S and Cy3-mRNA (Cy3 at position +20) to eliminate crosstalk from other dyes (**Fig. 3, A and B**) (*24*). *E*_FRET_ vs. time trajectories revealed a multi-frame ramp of *E*_FRET_ (mean lifetime ∼0.30 s) at the beginning of the FRET events before reaching the maximal *E*_FRET_ (∼0.48) on *RPL30* mRNA (**Fig. 3, C to H**). The ramp was more evident with *PMA1* mRNA (mean lifetime ∼0.58 s; **fig. S6, A and B**). Similar results were obtained at an increased movie frame rate (30 fps; **fig. S6, C to F**), indicating that the observed ramp was not an artifact of slow data acquisition. The mean ramp lifetime was similar to the scanning time measured above for *RPL30*, suggesting that the 43S landing site on *RPL30* corresponded to a distance that would allow rapid appearance of mRNA–uS5 FRET; whereas on *PMA1*, the ramp time was a fraction of the scanning time, consistent with 43S landing near the 5′ end of the mRNA and thus starting scanning from a position too distant for mRNA–uS5 FRET (**fig. S7A**).

**Fig. 3.**
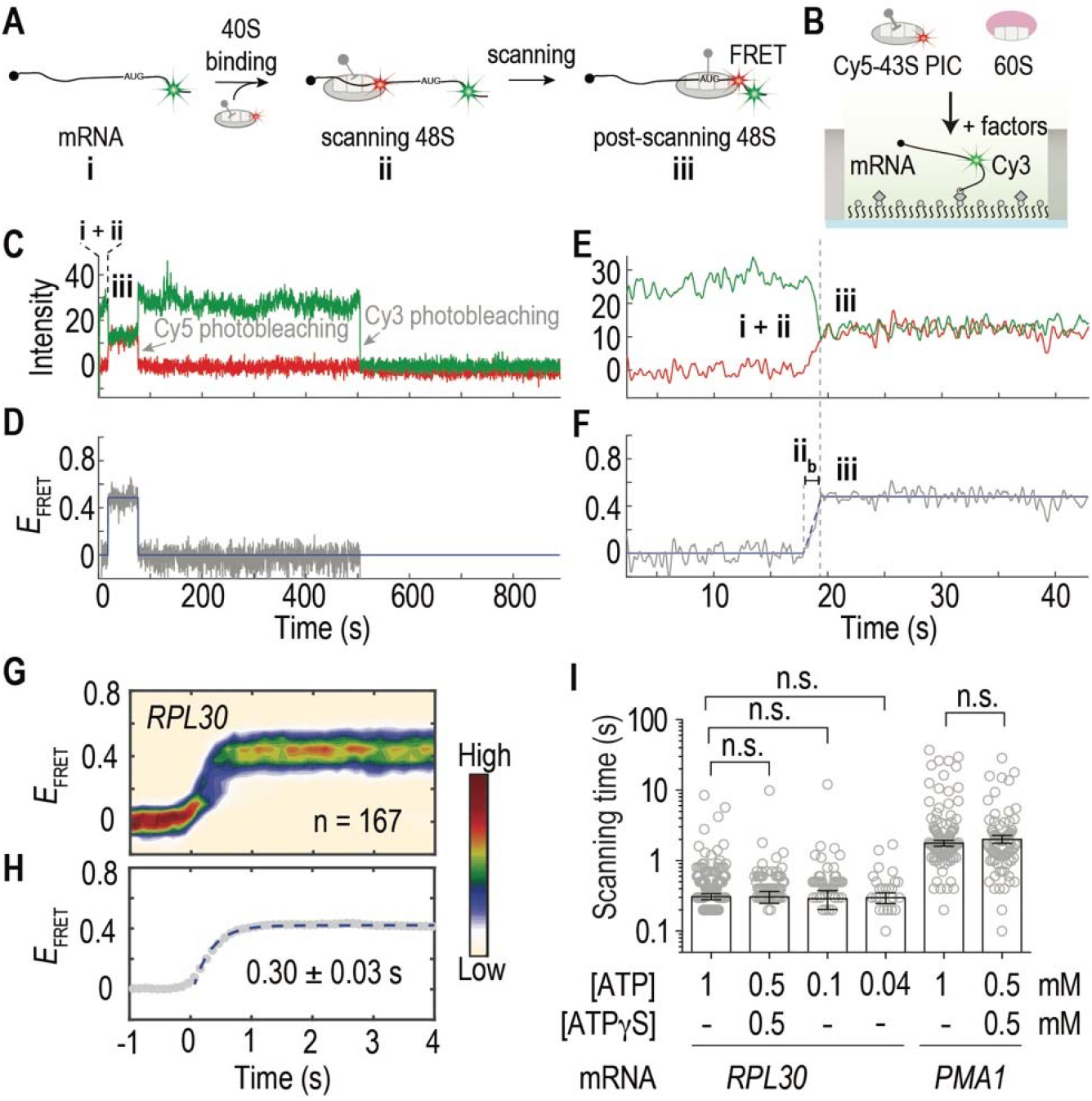
Scanning is globally 5′ to 3′ unidirectional and independent of excess eIF4A–ATP hydrolysis. (**A**) Schematic of assay to track scanning by two-color uS5-Cy5-40S to Cy3-mRNA FRET. (**B**) Assay setup. Capped-mRNA was labeled at +20 position with Cy3, tethered in ZMWs and pretreated with factors (including eIF4 proteins). uS5-Cy5-40S ribosomes were delivered along with other components to the ZMWs to start the reaction. (**C**) Example fluorescence and (**D**) *E*_FRET_ vs. time trajectory showing uS5-Cy5-40S binding to Cy3-mRNA, resulting in Cy3–Cy5 FRET. (**E**) and (**F**) show an expanded view at the beginning of the FRET event as in (**C**) and (**D**), which revealed a ramp of the *E*_FRET_ (state **ii**_**b**_). (**G**) All the *E*_FRET_ vs. time trajectories from experiments with *RPL30* mRNA were post-synchronized with the beginning of the FRET ramp set at time 0. (**H**) The post-synchronized *E*_FRET_ trajectories were averaged and fit to a single-exponential equation to estimate the mean FRET ramp time (± 95% CI). (**I**) Observed scanning times (open circles) measured under varying ATP conditions. Bars and errors represent mean scanning times ± 95% CI. (n.s., not significant, *P* > 0.4; two-tailed unpaired Student’s *t*-test). Data also shown in **fig. S6N**.

A small fraction of the FRET traces (∼11% for *RPL30* and ∼14% for *PMA1*) displayed *E*_FRET_ transitions between the maximum level and a lower non-zero state (**fig. S7, B and C**). Thus, 40S scanning is globally 5′ to 3′ unidirectional, with some occasional short-lived backward (3′ to 5′) movements (mean lifetime ∼2 s, **fig. S7D**), as suggested by lower mRNA–uS5 FRET levels (more below).

### Scanning does not require multiple ATP hydrolysis cycles on eIF4A

eIF4A is thought to drive scanning through rounds of ATP hydrolysis (*1, 20, 25*). Decreasing eIF4A concentration from 6 to 0.6 μM led to substantially slower appearance of the Cy3-mRNA to Cy5-uS5 FRET with *PMA1* mRNA (**fig. S6M**), consistent with the eIF4A requirement for 43S recruitment (**Fig. 1E**). However, the *E*_FRET_ ramp was largely unaffected (**fig. S6, G to J**), indicating that once a 43S PIC is loaded onto a mRNA, its scanning time is independent of eIF4A concentration. The scanning time determined in the four-color assay (**Fig. 2A**) remained similar when sub-saturating ATP concentrations (0.1 or 0.04 mM) (*26*) or a mixture of 0.5 mM ATP and ATPγS were used in place of 1 mM ATP (**Fig. 3I**). These results suggest that 43S scanning does not require multiple cycles of ATP hydrolysis by eIF4A. At a lower temperature (20°C), scanning was more heterogeneous on *PMA1* mRNA, with more ribosomes taking a slower path to the final FRET state and the mean ramp lifetime increased to ∼1.4 s (**fig. S6, K and L**).

### 5′UTR secondary structures modulate scanning dynamics

To determine how scanning is regulated by 5′UTR secondary structures, we engineered *RPL30* mRNA to contain a stable hairpin stem-loop (SL_1_ in mRNA-SL_1_-46, **Fig. 4A**, ΔG_mfe_ = −52.1 kcal/mol at 30ºC (*27*)). With SL_1_ located 60 nt from the 5′end and 46 nt upstream from the start codon, the two-color mRNA-uS5 FRET revealed a similar *E*_FRET_ ramp as observed for *RPL30*, without significant stalling of the scanning ribosomes (**fig. S8A**). In the four-color scanning assay (**Fig. 2A**), this hairpin structure had moderate effects on 43S recruitment and 60S joining (**fig. S9, A, C and D**). The mean scanning time (∼1.5 s) corresponded to a speed of 101 ± 13 nt•s^-1^ (**fig. S9B**), ∼30% slower than that determined above for scanning on 5′UTRs without highly stable structures. The stable SL_1_ in the middle of the 5′UTR thus slowed the scanning ribosomes, albeit slightly.

**Fig. 4.**
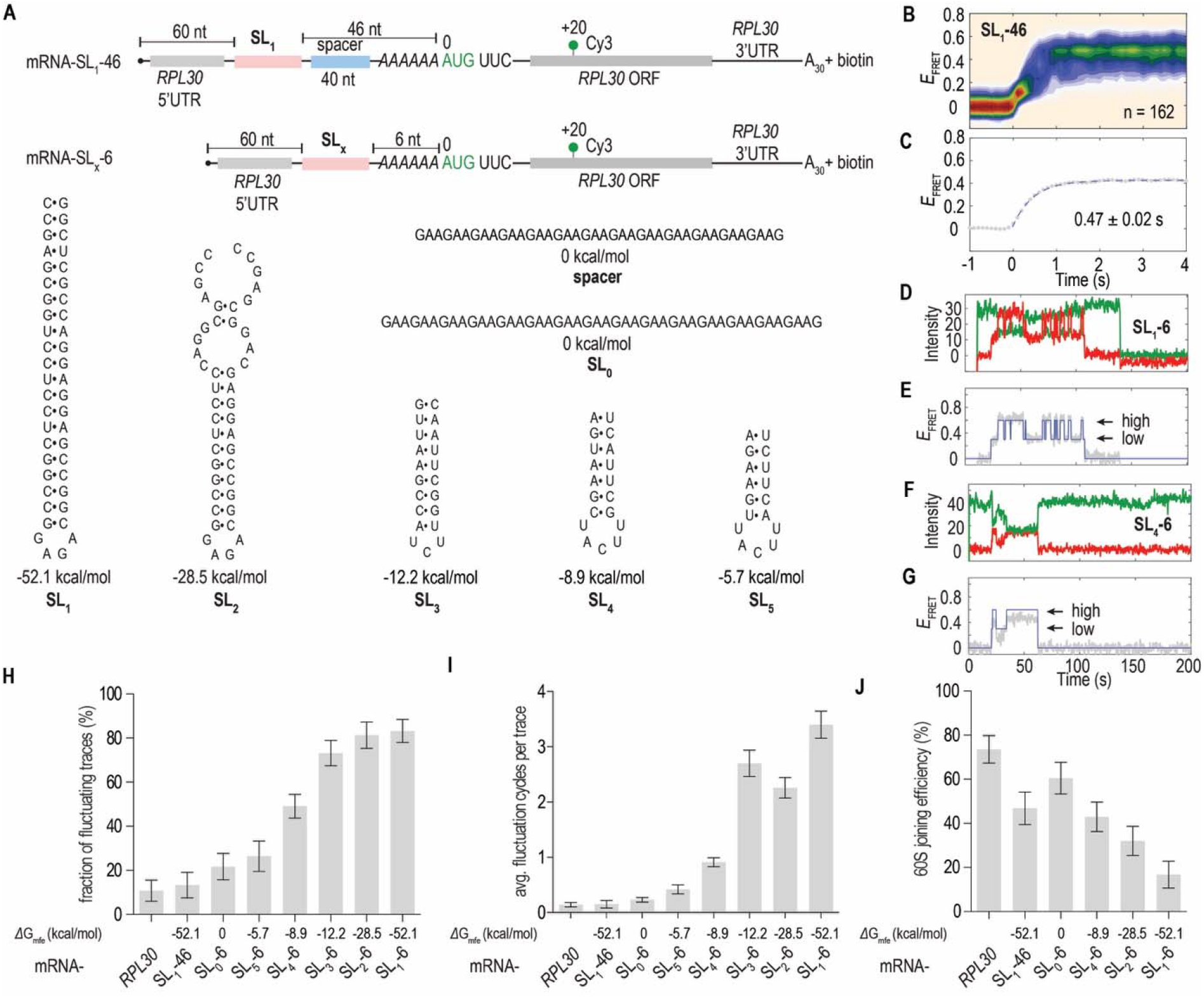
Secondary structures in 5′UTR modulate scanning dynamics. (**A**) mRNAs with varying structural elements in the 5′UTR. (**B**) Post-synchronized two-color *E*_FRET_ trajectories for mRNA-SL_1_-46 and (**C**) the average trajectory fit to a single-exponential equation to estimate the mean FRET ramp time (± 95% CI). Sample trace and *E*_FRET_ trajectory for mRNA-SL_1_-6 (**D** and **E**) or mRNA-SL_4_-6 (**F** and **G**) showing mRNA–uS5 FRET fluctuations. (**H**) Fraction of traces displayed FRET fluctuations and (**I**) average fluctuation cycles per trace increased with, while (**J**) 60S joining efficiencies (i.e. fraction of 48S PICs joined by 60S) decreased with the AUG-proximal hairpin stability. Data were shown as mean ± 95% CI and also in **fig. S9, D and I**.

Previous analyses in yeast suggested larger translation inhibition effects of 5’UTR secondary structures located proximal rather than distal to the start site (*12*). We thus moved SL_1_ closer to the AUG codon (mRNA-SL_1_-6, **Fig. 4A**). Cy3-mRNA to uS5-Cy5 FRET also rapidly arrived at a high FRET state, but ∼80% of the FRET traces displayed discrete fluctuations (average ∼3 cycles) between the high (∼0.47) and a low (∼0.18) FRET state (**Fig.4, D, E, H and I, fig. S8A**). The high FRET state correlated with the 43S at the AUG site, consistent with 43S scanning through the hairpin; the fluctuation to the low FRET state (with abrupt state transitions, **fig. S9J**) reflected the backward (3′ to 5′) movement of the ribosome induced by the hairpin sequence; and the following transition to the high FRET state indicated “rescanning” by the ribosome to the AUG site. In agreement with this interpretation, an unlabeled DNA oligonucleotide blocking the start site suppressed the high FRET state with mRNA-SL_1_-6 (**fig. S8**). Replacing GTP with the non-hydrolyzable analog GDPCP did not alter the fluctuation dynamics (**fig. S9, E, F and I**), indicating that the fluctuations can occur prior to GTP hydrolysis by eIF2.

When SL_1_ was replaced with lower-stability RNA structures (−28.5 kcal/mol and lower, **Fig. 4A**), similar FRET fluctuations were observed, but with reduced frequencies (**Fig. 4, F to I, fig. S9, G to I, and fig. S10**); fluctuations were not affected by the presence of Ded1p. The low FRET state lifetimes increased with hairpin stability (**fig. S9, G and I**), consistent with increased free energies to rescan through the hairpins. Lowering the reaction temperature to 20ºC led to a >twofold increase in the low FRET state lifetimes with mRNA-SL_1_-6 (**fig. S9, E and I**), consistent with increased hairpin stability at lower temperature. Reducing ATP concentration from 1 to 0.1 mM resulted in a ∼1.7-fold increase in the low FRET state lifetimes, without changing the high FRET state lifetimes (**fig. S9, E, F and I**). Decreasing eIF4A concentration from 6 to 0.6 μM also prolonged the low FRET state lifetimes (**fig. S9, E, F and I**). Thus, multiple cycles of ATP hydrolysis and rebinding of eIF4A promote the resumption of 40S scanning towards the 3′ direction during the fluctuations induced by stable hairpins.

With more stable hairpin structures, fewer ribosomes stabilized at the high FRET state, as shown by the less populated high FRET state (**fig. S10**) and substantially reduced high FRET state lifetimes (**fig. S9, H and I**). Consistently, both the 60S joining efficiencies and rates were reduced as a function of hairpin stability in the four-color scanning assay (**Fig. 4J, fig. S9, C and D**).

### Downstream hairpin structure stimulates near-cognate initiation

Initiation can occur at near-cognate start codons (e.g. CUG), but at relatively low levels and often stimulated by downstream hairpin structures (*28, 29*). We deployed a single-molecule assay (**Fig. 5, A to D**) to distinguish initiation at the CUG site (within 5′UTR) and the AUG site, by FRET between Cy3-60S and the first elongator tRNA of the two reading frames (Cy5.5-Phe-tRNA^Phe^ for AUG and Cy5-Lys-tRNA^Lys^ for CUG) (**fig. S11**). A downstream stable SL_1_ hairpin led to a fourfold increase in the fraction of ribosomes initiated at the CUG codon compared with SL_0_ (∼48.4% for CUG-SL_1_-AUG vs. ∼11.9% for CUG-SL_0_-AUG mRNAs, **Fig. 5E**).

**Fig. 5.**
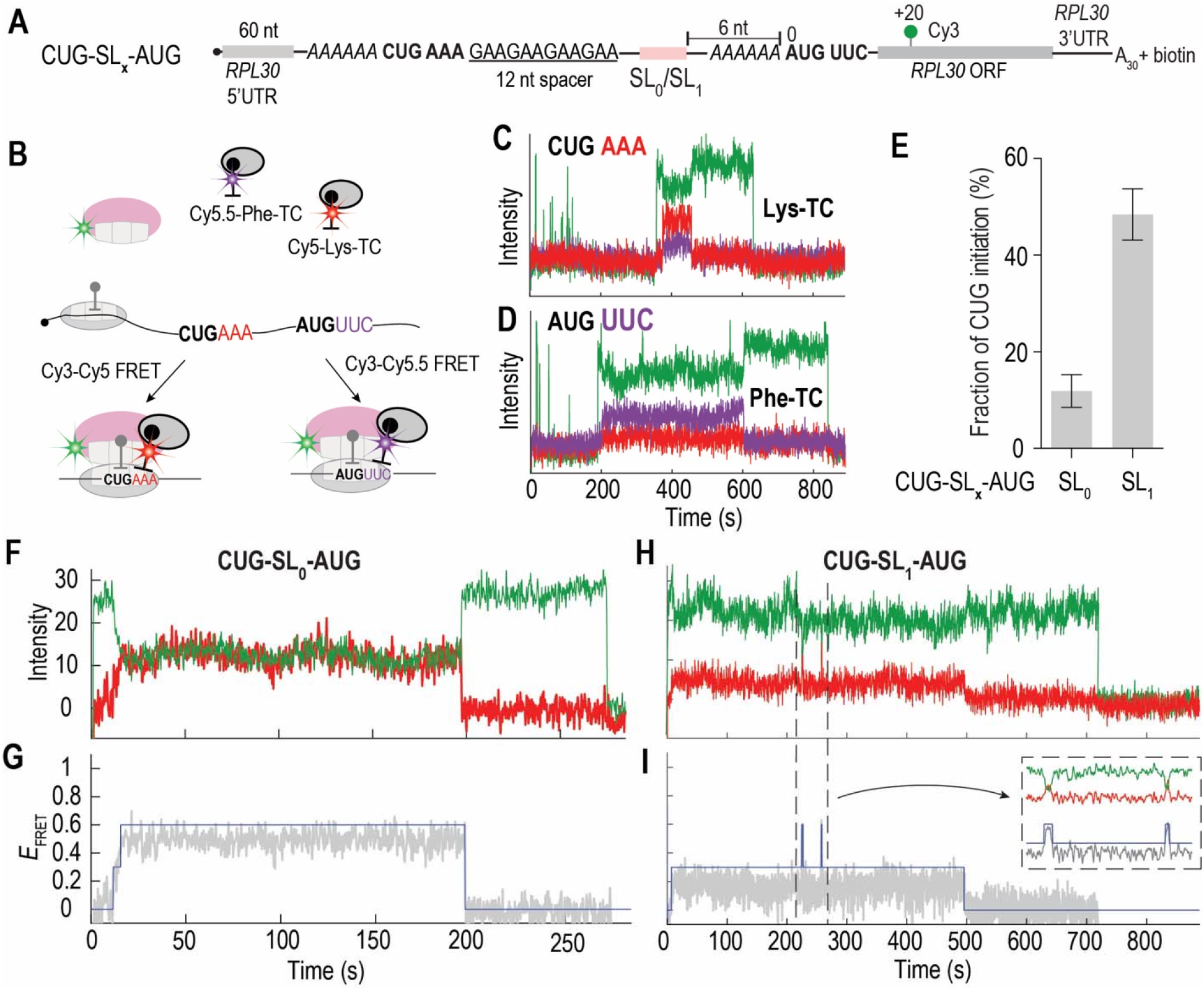
Secondary structures stimulate upstream near-cognate codon initiation. (**A**) mRNAs with an upstream CUG codon and SL_0_ or SL_1_ sequence in the 5′UTR. (**B**) Single-molecule assay setup and sample traces (**C** and **D**) to detect CUG vs. AUG initiation by 60S–elongator tRNA FRET (see **fig. S11**). (**E**) Fraction of CUG initiation out of total initiation events at both CUG and AUG sites in the presence (n = 337) or absence (n = 352) of a stable hairpin structure (mean ± 95% CI). Sample two-color mRNA–uS5 FRET traces and *E*_FRET_ trajectories for mRNAs CUG-SL_0_-AUG (**F** and **G**) and CUG-SL_1_-AUG (**H** and **I**).

Next, we tested near-cognate initiation in our two-color scanning assay. With CUG-SL_0_- AUG mRNA, ∼12% of the mRNA–uS5 FRET trajectories showed only a long-lived (mean lifetime ∼124 s) low FRET state (*E*_FRET_ ∼0.17, **fig. S12, A to C**), presumably due to the scanning ribosomes initiating at the CUG site during the first encounter. The rest of the trajectories displayed a high FRET state (*E*_FRET_ ∼0.50), either without fluctuation (∼64%) or fluctuated to the low FRET state but eventually stayed in a long-lived high FRET state (∼24%, and lifetime with a dominant slow phase at 127 ± 2 s) (**Fig. 5, F and G, fig. S12, C and D**). These likely reflected the ribosomes that bypassed the upstream CUG codon and initiated at the AUG site.

With downstream SL_1_, the fraction of low FRET state-only trajectories increased to ∼21%, consistent with a higher energy barrier to unfold the hairpin by the scanning ribosome. However, these were insufficient to account for the ∼48.4% CUG initiation with this mRNA (**Fig. 5E**). Most trajectories (∼77%) displayed FRET state fluctuations after reaching the high FRET state (**Fig. 5, H and I**), with the low FRET state lifetime being ∼3-fold longer than those observed above for mRNA-SL_1_-6 (**fig. S12, E and F**), suppressing the occupancy of the high FRET state (**fig. S12C**). These trajectories plausibly correspond to scanning 40S that bypassed the CUG codon on first passage and reached the AUG site, where the AUG site-proximal hairpin induced ribosome fluctuations to increase the number of times a ribosome samples the upstream CUG codon, thus increasing CUG initiation. Consistent with a delay in initiation caused by ribosomal fluctuations on the mRNA, 60S binding events were ∼fivefold slower compared to CUG-SL_0_-AUG at both the CUG and AUG sites (**fig. S12G**). Moving the upstream CUG codon and SL_1_ hairpin 40 nt away from the AUG site (**fig. S12G**), which abolishes the ribosome fluctuations, decreased the CUG initiation efficiency back to ∼11% (**fig. S12H**).

## Discussion

Despite longstanding evidence for ribosomal scanning in canonical eukaryotic translation initiation (*1*–*7*), the dynamic scanning motion of the 43S has not been directly observed. Our single-molecule fluorescence approaches quantitatively monitor translation initiation directly as it unfolds from 43S–mRNA binding through scanning and 60S subunit joining, providing a foundation to understand its regulation (**Fig. 6**).

**Fig. 6.**
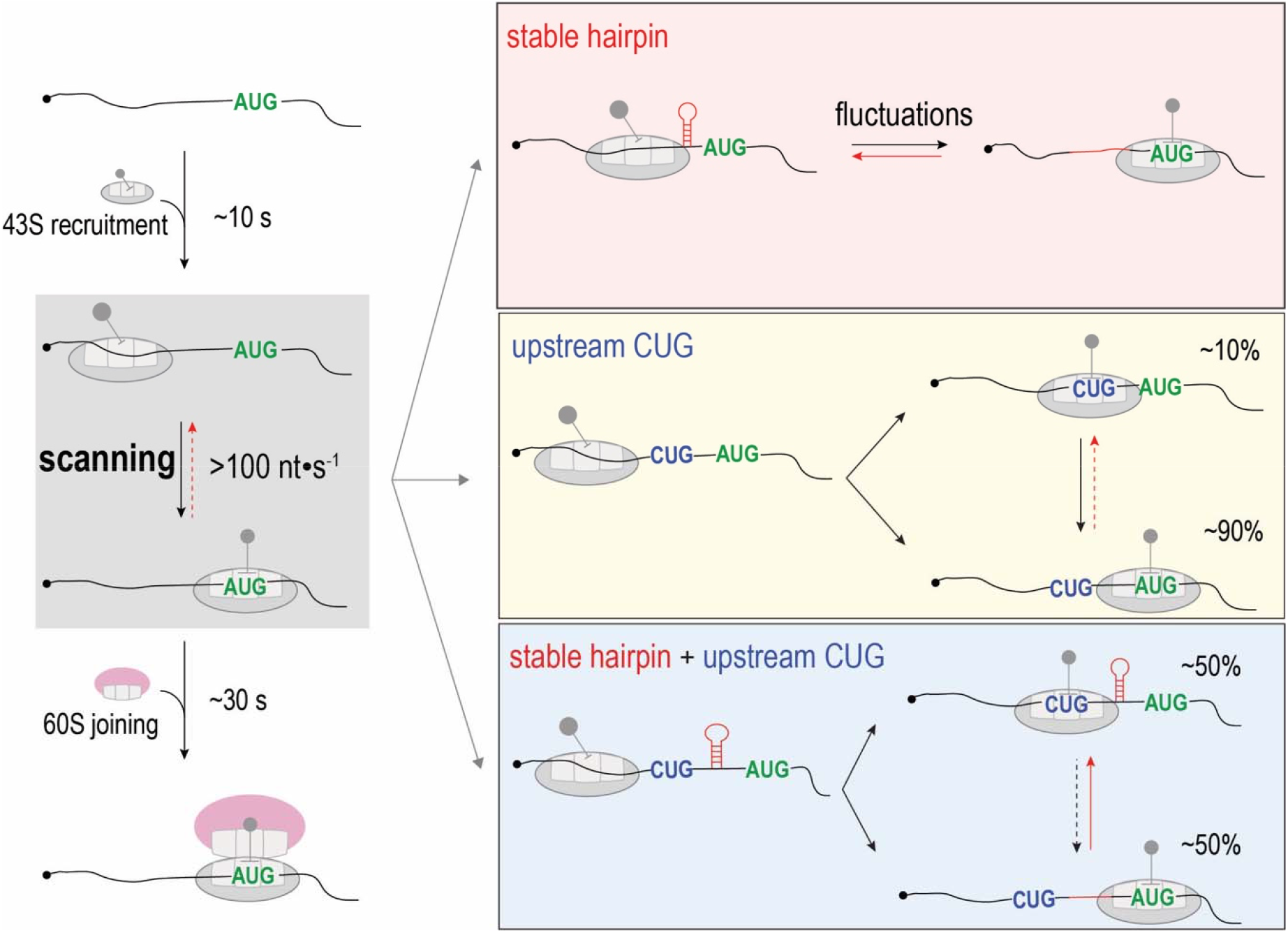
A quantitative model of eukaryotic translation initiation. Rapid 43S PIC recruitment to 5′ capped-mRNA is catalyzed by eIF4F proteins, especially ATP hydrolysis by eIF4A. Once loaded, the 43S PIC scans the 5′UTR with a globally 5′ to 3′ direction and a speed >100 nt•s^-1^, independent of multiple rounds of ATP hydrolysis by eIF4A. Upon arriving to the start codon, the ribosome complex undergoes slow rearrangements before 60S can join. The scanning 43S PIC can unfold stable 5′UTR hairpin structure and reach the start site. When the hairpin is adjacent to the start codon, it induces backward (3′ to 5′) movement of the 43S PIC; when an upstream near-cognate (CUG) codon is present, the AUG-proximal hairpin stimulates initiation at the near-cognate codon largely by increasing the number of times the 43S PIC encounters and resides at the near-cognate codon.

Efficient 43S recruitment requires the preparation of mRNAs by eIF4F proteins, in particular eIF4A and its ATP hydrolysis (**Fig. 1E** and **fig. S2**), prerhaps through the unwinding of the mRNA structures either locally around the 40S landing site or globally throughout the entire sequence (*25*), or mechanically aid mRNA loading within the 40S mRNA binding cleft (*30*). This could rationalize the high abundance of eIF4A in cells (> 10 μM) and impaired global translation with reduced eIF4A concentrations (*3, 31*). By contrast, the more effective RNA helicase Ded1p (*19*) was unable to promote 40S-mRNA binding in the absence of eIF4A, implying that the RNA helicase activity alone is insufficient to drive initiation. Thus, eIF4A is an indispensable constituent of the eIF4F-mRNA intermediate that is compatible with rapid 43S binding (*25, 32*) and may aid 43S PIC recruitment through remodeling of 40S conformations (*25, 33*).

Once loaded onto the mRNA, 43S scanning through yeast 5′UTRs is rapid (>100 nt•s^-1^) and may be coupled to loading. The 43S PIC landing site on the mRNA is near the 5′ end, and once engaged with the mRNA, it is critical for the 40S subunit to commit to scanning and subsequent initiation, as failures would lead to stable stalled complexes (**fig. S5, E and G**), sequestering the 40S subunits and bound eIFs on an mRNA. Fast scanning represents a technical challenge to our single-molecule FRET approach (30 ms temporal resolution) for obtaining nucleotide-resolution information on its directionality. Nonetheless, our results indicate that scanning is globally 5′ to 3′ unidirectional, with a finite low propensity to backward movement (**Fig. 3**). The scanning motion does not rely on multiple rounds of eIF4A-ATP hydrolysis and is independent of Ded1p. What governs the scanning speed and directionality remains to be established, but the data presented here cannot rule out a biased diffusional mechanism over the relatively short 5′UTR distances that must be scanned by an initiating yeast ribosome (*34*). More heterogeneous trajectories of scanning were observed at lower temperature (**fig. S6K**), suggesting that scanning is modulated by energy barriers, e.g. RNA structures in the 5′UTR. Scanning may be directly coupled to the energy expenditure by eIF4A during 43S PIC recruitment, and 5′ to 3′ directionality may be linked to the 43S–mRNA binding energy, and contributions of eIF3 and eIF4F proteins bound at the mRNA exit site (*32*).

Scanning is not rate-limiting in initiation under normal conditions with unstructured 5′UTRs. After rapid 43S scanning to the start site, a relatively long delay (post-scanning state, ∼30 s to minutes) to 60S joining occurs (**fig. S4, B and C**), which limits the overall initiation rate. During this time period, the ribosomal complex likely undergoes conformational and compositional rearrangements, leading to stable initiator tRNA anticodon-start codon interactions before commitment to the final stages of initiation. eIF2 must hydrolyze GTP and dissociate from the ribosome to enable eIF5B recruitment and subsequent 60S joining (*35*), while eIF3 configurations have been shown to remodel upon start codon recognition (*36, 37*). The presence of Ded1p shortened this delay by ∼2-fold (**fig. S4, B and C**), plausibly through its interplay with eIF4F proteins, the 40S ribosome and the mRNA (*18, 19, 38*–*40*). This is consistent with prior evidence that its human homolog, DDX3, assists 60S joining to assemble functional 80S ribosomes (*41*).

5′UTR secondary structures play a surprising role in scanning. Here we demonstrate that scanning ribosomes readily melt mRNA hairpin structures up to -52.1 kcal/mol in stability in the 5′UTR upon first encounter to reach the start site rapidly (**Fig. 4**). However, when the hairpin is located adjacent to the start codon, the ribosome remains at the start site for only a short period (∼3 s, **fig. S9I**); subsequent backwards motion (3′ to 5′) and fluctuations between the upstream position and the start site impair 60S joining. The frequency of such fluctuations plateaued with hairpin stabilities above -12.2 kcal/mol (**Fig. 4, H and I**). This provides a rationale for the underrepresented (∼1%) secondary structures with stability above -12.2 kcal/mol in a 50-nt window in yeast 5′UTRs (*13*), and explains why secondary structures immediately upstream of the start site (positions -40 to -1) are selected against in yeast (*42*).

Fluctuations are induced by hairpin structures that are unfolded within the 40S subunit mRNA binding channel. The hairpin-induced 3′ to 5′ movement of the 40S occurs prior to GTP hydrolysis by eIF2 (**fig. S9, E, F and I**) during the long waiting period for 60S subunit joining. Reverse movement of the 40S subunit may be the result of hairpin refolding on the ribosome near the mRNA exit channel, leading to steric clashes with eIF3 (and perhaps eIF4F) components (*32, 43*) and thus biasing the 40S subunit to positions upstream (5′) of the hairpin. Moving the hairpin further upstream from the AUG diminished this regulatory effect (**Fig. 4, A to C**); in this case the ribosomal occupancy with hairpin unfolded is transient (< 0.5 s) due to the rapid rate (>100 nt•s^-1^) of scanning to the downstream start codon, leading to a low probability of hairpin refolding on the ribosome. The high-to-low uS5-mRNA FRET transitions were nearly instantaneous (**fig. S9J**), suggesting abrupt 40S backward movement to upstream of the hairpin coupled to rapid two-state hairpin refolding and thus reduced 40S occupancies within the hairpin sequence. Consistent with this notion, re-analysis of the human 40S ribosome profiling data (*44*) (too sparse data for yeast due to generally unstructured 5′UTRs) revealed reduced 40S occupancies within a 50-nt window immediately upstream of the start codon, when that region of the mRNA has a high propensity (ΔG_mfe_ beyond -20 kcal/mol) for hairpin structure formation (**fig. S13**).

Scanning 40S subunits in our assay largely bypass (∼90%) upstream near-cognate CUG codons, even within the optimal sequence context (**Fig. 5E**), consistent with the low in vivo efficiencies of initiation at near-cognate start codons (*45, 46*). Stable hairpin structures downstream of the CUG codon enhanced initiation at near-cognate codons by two mechanisms. First, the increase in activation energy required to unwind the hairpin during forward scanning moderately increases the fraction of ribosomes that settle and initiate at the near-cognate CUG upon first encounter (**fig. S12, A and B**) (*28*). Second, when the 5’UTR hairpin is proximal to the AUG codon, scanning ribosomes that reach the AUG site are driven backwards and fluctuate (**Fig. 5H**), enhancing CUG initiation efficiency by increasing the number of times a ribosome encounters and resides at the CUG codon during the delayed period of initiation (**Fig. 5E**).

Our study establishes a quantitative mechanistic framework of eukaryotic translation initiation (**Fig. 6**), and the dynamic mechanisms unraveled here will guide future efforts on human translation to understand translational control in health and disease.

## Acknowledgements

We thank Rosslyn Grosely, Michael Lawson and Alex Johnson for comments, Alexey Petrov and Zheren Ou for help with ebFRET data analysis pipeline, and Israel Fernández and Puglisi lab members, especially Christopher Lapointe and Rosslyn Grosely, for discussion.

## Author contributions

Conceptualization: JW, JDP

Methodology: JW, CA, BSS, JB, TED, JDP

Investigation: JW, CA, BSS, JB

Visualization: JW, JB

Supervision: TED, JDP

Writing – original draft: JW, JDP

Writing – review & editing: JW, CA, BSS, JB, TED, JDP

## Data and materials availability

All data is available in the manuscript or the supplementary materials. Expression constructs and yeast strains are available upon request.

## List of Supplementary Materials

Materials and Methods

Figs. S1-S13

Table S1

References (47-57)

## References

1. A. G. Hinnebusch, The scanning mechanism of eukaryotic translation initiation. Annu. Rev. Biochem. 83, 779–812 (2014).

2. C. E. Aitken, J. R. Lorsch, A mechanistic overview of translation initiation in eukaryotes. Nat. Struct. Mol. Biol. 19, 568–576 (2012).

3. W. C. Merrick, G. D. Pavitt, Protein synthesis initiation in eukaryotic cells. Cold Spring Harb. Perspect. Biol. 10, a033092 (2018).

4. M. Sokabe, C. S. Fraser, Toward a kinetic understanding of eukaryotic translation. Cold Spring Harb. Perspect. Biol. 11, a032706 (2019).

5. R. J. Jackson, C. U. T. Hellen, T. V. Pestova, The mechanism of eukaryotic translation initiation and principles of its regulation. Nat. Rev. Mol. Cell Biol. 11, 113–127 (2010).

6. A. G. Hinnebusch, Structural insights into the mechanism of scanning and start codon recognition in eukaryotic translation initiation. Trends Biochem. Sci. 42, 589–611 (2017).

7. M. Kozak, How do eucaryotic ribosomes select initiation regions in messenger RNA? Cell. 15, 1109–1123 (1978).

8. M. Siwiak, P. Zielenkiewicz, A comprehensive, quantitative, and genome-wide model of translation. PLOS Comput. Biol. 6, 1000865 (2010).

9. R. D. Palmiter, Quantitation of parameters that determine the rate of ovalbumin synthesis. Cell. 4, 189–197 (1975).

10. P. Shah, Y. Ding, M. Niemczyk, G. Kudla, J. B. Plotkin, Rate-limiting steps in yeast protein translation. Cell. 153, 1589–1601 (2013).

11. K. Leppek, R. Das, M. Barna, Functional 5′ UTR mRNA structures in eukaryotic translation regulation and how to find them. Nat. Rev. Mol. Cell Biol. 19, 158–174 (2018).

12. J. T. Cuperus, B. Groves, A. Kuchina, A. B. Rosenberg, N. Jojic, S. Fields, G. Seelig, Deep learning of the regulatory grammar of yeast 5′ untranslated regions from 500,000 random sequences. Genome Res. 27, 2015–2024 (2017).

13. M. Ringné, M. Krogh, Folding free energies of 5′-UTRs impact post-transcriptional regulation on a genomic scale in yeast. PLOS Comput. Biol. 1, e72 (2005).

14. J. R. Babendure, J. L. Babendure, J.-H. Ding, R. Y. Tsien, Control of mammalian translation by mRNA structuren near caps. RNA. 12, 851–861 (2006).

15. R. Y. Chuang, P. L. Weaver, Z. Liu, T. H. Chang, Requirements of the DEAD-box protein Ded1p for messenger RNA translation. Science. 275, 1468–1471 (1997).

16. N. D. Sen, F. Zhou, N. T. Ingolia, A. G. Hinnebusch, Genome-wide analysis of translational efficiency reveals distinct but overlapping functions of yeast DEAD-box RNA helicases Ded1 and eIF4A. Genome Res. 25, 1196–1205 (2015).

17. D. Sharma, E. Jankowsky, The Ded1/DDX3 subfamily of DEAD-box RNA helicases. Crit. Rev. Biochem. Mol. Biol. 49, 343–360 (2014).

18. A. Hilliker, Z. Gao, E. Jankowsky, R. Parker, The DEAD-Box protein Ded1 modulates translation by the formation and resolution of an eIF4F-mRNA complex. Mol. Cell. 43, 962–972 (2011).

19. Z. Gao, A. A. Putnam, H. A. Bowers, U.-P. Guenther, X. Ye, A. Kindsfather, A. K. Hilliker, E. Jankowsky, Coupling between the DEAD-box RNA helicases Ded1p and eIF4A. Elife. 5, e16408 (2016).

20. W. C. Merrick, eIF4F: a retrospective. J. Biol. Chem. 290, 24091–24099 (2015).

21. J. Wang, A. G. Johnson, C. P. Lapointe, J. Choi, A. Prabhakar, D.-H. Chen, A. N. Petrov, J. D. Puglisi, eIF5B gates the transition from translation initiation to elongation. Nature. 573, 605–608 (2019).

22. J. Wang, J. Wang, B.-S. Shin, J.-R. Kim, T. E. Dever, J. D. Puglisi, I. S. Fernández, Structural basis for the transition from translation initiation to elongation by an 80S-eIF5B complex. Nat. Commun. 11, 5003 (2020).

23. T. Tuller, E. Ruppin, M. Kupiec, Properties of untranslated regions of the S. cerevisiae genome. BMC Genomics. 10, 391 (2009).

24. J. Chen, R. V. Dalal, A. N. Petrov, A. Tsai, S. E. O’Leary, K. Chapin, J. Cheng, M. Ewan, P.-L. Hsiung, P. Lundquist, S. W. Turner, D. R. Hsu, J. D. Puglisi, High-throughput platform for real-time monitoring of biological processes by multicolor single-molecule fluorescence. Proc. Natl. Acad. Sci. 111, 664–669 (2014).

25. P. Yourik, C. E. Aitken, F. Zhou, N. Gupta, A. G. Hinnebusch, J. R. Lorsch, Yeast eIF4A enhances recruitment of mRNAs regardless of their structural complexity. Elife. 6, e31476 (2017).

26. J. R. Lorsch, D. Herschlag, The DEAD box protein eIF4A. 1. A minimal kinetic and thermodynamic framework reveals coupled binding of RNA and nucleotide. Biochemistry. 37, 2180–2193 (1998).

27. M. Zuker, Mfold web server for nucleic acid folding and hybridization prediction. Nucleic Acids Res. 31, 3406–3415 (2003).

28. M. Kozak, Downstream secondary structure facilitates recognition of initiator codons by eukaryotic ribosomes. Proc. Nail. Acad. Sci. USA. 87, 8301–8305 (1990).

29. M. G. Kearse, J. E. Wilusz, Non-AUG translation: a new start for protein synthesis in eukaryotes. Genes Dev. 31, 1717–1731 (2017).

30. M. Sokabe, C. S. Fraser, A helicase-independent activity of eIF4A in promoting mRNA recruitment to the human ribosome. Proc. Nail. Acad. Sci. USA. 114, 6304–6309 (2017).

31. H. Firczuk, S. Kannambath, J. Pahle, A. Claydon, R. Beynon, J. Duncan, H. Westerhoff, P. Mendes, J. E. McCarthy, An in vivo control map for the eukaryotic mRNA translation machinery. Mol. Syst. Biol. 9, 635 (2013).

32. J. B. Querido, M. Sokabe, S. Kraatz, Y. Gordiyenko, J. M. Skehel, C. S. Fraser, V. Ramakrishnan, Structure of a human 48S translational initiation complex. Science. 369, 1220–1227 (2020).

33. Y. Gu, Y. Mao, L. Jia, L. Dong, S.-B. Qian, Bi-directional ribosome scanning controls the stringency of start codon selection. Nat. Commun. 12, 6604 (2021).

34. K. S. Vassilenko, O. M. Alekhina, S. E. Dmitriev, I. N. Shatsky, A. S. Spirin, Unidirectional constant rate motion of the ribosomal scanning particle during eukaryotic translation initiation. Nucleic Acids Res. 39, 5555–5567 (2011).

35. M. A. Algire, D. Maag, J. R. Lorsch, Pi release from eIF2, not GTP hydrolysis, is the step controlled by start-site selection during eukaryotic translation initiation. Mol. Cell. 20, 251–262 (2005).

36. J. L. Llácer, T. Hussain, J. Dong, L. Villamayor, Y. Gordiyenko, A. G. Hinnebusch, Large-scale movement of eIF3 domains during translation initiation modulate start codon selection. Nucleic Acids Res. 49, 11491–11511 (2021).

37. A. Simonetti, J. Brito Querido, A. G. Myasnikov, E. Mancera-Martinez, A. Renaud, L. Kuhn, Y. Hashem, eIF3 Peripheral Subunits Rearrangement after mRNA Binding and Start-Codon Recognition. Mol. Cell. 63, 206–217 (2016).

38. S. Gulay, N. Gupta, J. R. Lorsch, A. G. Hinnebusch, Distinct interactions of eIF4A and eIF4E with RNA helicase Ded1 stimulate translation in vivo. Elife. 9, e58243 (2020).

39. N. Gupta, J. R. Lorsch, A. G. Hinnebusch, Yeast Ded1 promotes 48S translation preinitiation complex assembly in an mRNA-specific and eIF4F-dependent manner. Elife. 7, e38892 (2018).

40. U. P. Guenther, D. E. Weinberg, M. M. Zubradt, F. A. Tedeschi, B. N. Stawicki, L. L. Zagore, G. A. Brar, D. D. Licatalosi, D. P. Bartel, J. S. Weissman, E. Jankowsky, The helicase Ded1p controls use of near-cognate translation initiation codons in 5′ UTRs. Nature. 559, 130–134 (2018).

41. R. Geissler, R. P. Golbik, S. E. Behrens, The DEAD-box helicase DDX3 supports the assembly of functional 80S ribosomes. Nucleic Acids Res. 40, 4998–5011 (2012).

42. A. Robbins-Pianka, M. D. Rice, M. P. Weir, A. Bateman, The mRNA landscape at yeast translation initiation sites. Bioinformatics. 26, 2651–2655 (2010).

43. J. L. Llácer, T. Hussain, L. Marler, C. E. Aitken, A. Thakur, J. R. Lorsch, A. G. Hinnebusch, V. Ramakrishnan, Conformational differences between open and closed states of the eukaryotic translation initiation complex. Mol. Cell. 59, 399–412 (2015).

44. J. Bohlen, K. Fenzl, G. Kramer, B. Bukau, A. A. Teleman, Selective 40S footprinting reveals cap-tethered ribosome scanning in human cells. Mol. Cell. 79, 561–574 (2020).

45. A. R. Eisenberg, A. L. Higdon, I. Hollerer, A. P. Fields, I. Jungreis, P. D. Diamond, M. Kellis, M. Jovanovic, G. A. Brar, Translation initiation site profiling reveals widespread synthesis of non-AUG-initiated protein isoforms in yeast. Cell Syst. 11, 145–160 (2020).

46. G. Loughran, A. V. Zhdanov, M. S. Mikhaylova, F. N. Rozov, P. N. Datskevich, S. I. Kovalchuk, M. V. Serebryakova, S. J. Kiniry, A. M. Michel, P. B. F. O’Connor, D. B. Papkovsky, J. F. Atkins, P. V. Baranov, I. N. Shatsky, D. E. Andreev, Unusually efficient CUG initiation of an overlapping reading frame in POLG mRNA yields novel protein POLGARF. Proc. Natl. Acad. Sci. U. S. A. 117, 24936–24946 (2020).

47. S. F. Mitchell, S. E. Walker, M. A. Algire, E. H. Park, A. G. Hinnebusch, J. R. Lorsch, The 5′-7-methylguanosine cap on eukaryotic mRNAs serves both to stimulate canonical translation initiation and to block an alternative pathway. Mol. Cell. 39, 950–962 (2010).

48. E. H. Park, S. E. Walker, J. M. Lee, S. Rothenburg, J. R. Lorsch, A. G. Hinnebusch, Multiple elements in the eIF4G1 N-terminus promote assembly of eIF4G1•PABP mRNPs in vivo. EMBO J. 30, 302–316 (2011).

49. A. Petrov, R. Grosely, J. Chen, S. E. O’Leary, J. D. Puglisi, Multiple parallel pathways of translation initiation on the CrPV IRES. Mol. Cell. 62, 92–103 (2016).

50. O. Puig, B. Rutz, B. B. M. Luukkonen, S. Kandels-Lewis, E. Bragado-Nilsson, B. Séraphin, New constructs and strategies for efficient PCRLbased gene manipulations in yeast. Yeast. 14, 1139–1146 (1998).

51. Z. Zhou, P. Cironi, A. J. Lin, Y. Xu, S. Hrvatin, D. E. Golan, P. A. Silver, C. T. Walsh, J. Yin, Genetically encoded short peptide tags for orthogonal labelling by Sfp and AcpS phosphopantetheinyl transferases. ACS Chem. Biol. 2, 337–346 (2007).

52. M. R. Stark, J. A. Pleiss, M. Deras, S. A. Scaringe, S. D. Rader, An RNA ligase-mediated method for the efficient creation of large, synthetic RNAs. RNA. 12, 2014–2019 (2006).

53. J. W. van de Meent, J. E. Bronson, C. H. Wiggins, R. L. Gonzalez, Empirical Bayes Methods Enable Advanced Population-Level Analyses of Single-Molecule FRET Experiments. Biophys. J. 106, 1327–1337 (2014).

54. S. Peng, R. Sun, W. Wang, C. Chen, Single-molecule photoactivation FRET: a general and easy-to-implement approach to break the concentration barrier. Angew. Chemie. 129, 6986–6989 (2017).

55. E. F. Pettersen, T. D. Goddard, C. C. Huang, E. C. Meng, G. S. Couch, T. I. Croll, J. H. Morris, T. E. Ferrin, UCSF ChimeraX: Structure visualization for researchers, educators, and developers. Protein Sci. 30, 70–82 (2021).

56. R. Buschauer, Y. Matsuo, T. Sugiyama, Y. H. Chen, N. Alhusaini, T. Sweet, K. Ikeuchi, J. Cheng, Y. Matsuki, R. Nobuta, A. Gilmozzi, O. Berninghausen, P. Tesina, T. Becker, J. Coller, T. Inada, R. Beckmann, The Ccr4-Not complex monitors the translating ribosome for codon optimality. Science. 368, eaay6912 (2020).

57. A. R. Gruber, R. Lorenz, S. H. Bernhart, R. Neuböck, I. L. Hofacker, The Vienna RNA Websuite. Nucleic Acids Res. 36, W70–W74 (2008).

