## Supplementary Materials for "Rapid 40S scanning and its regulation by mRNA structure during eukaryotic translation initiation"

Jinfan Wang *et al.*

**This file includes:**

Materials and Methods

Figs. S1 to S13

Table S1

References (47-57)

**Materials and Methods**

**1. Yeast translation factors**

Yeast *Saccharomyces cerevisiae* eIFs 1, 1A, 2, 3, 4A, 4B, 4E, 4G, 5 and 5B, and eEFs 1A and 1Bα were prepared according to previously established methods (*21*). To express Pab1p or Hcr1p, the plasmid pTYB2-Pab1p (Addgene #37234) or pTYB2-HCR1 (Addgene #37233) were transformed into *Escherichia coli* Rosetta2 (DE3) cells (Novagen), and the proteins were purified as previously described (*47*, *48*).

Full-length *DED1* gene was PCR-amplified from yeast genomic DNA and cloned into a pET28c plasmid for the expression of a recombinant 6×His-MBP-TEV protease site-Ded1p fusion protein. The resulting plasmid was transformed into Rosetta2 (DE3) cells and overexpressed by induction at an OD_600nm_ = 0.5 with 1 mM IPTG at 16ºC for 16 hours. Cells were pelleted by centrifugation at 5000 × g for 12 min, resuspended in 30 mL of lysis buffer (50 mM HEPES-KOH pH 7.5, 300 mM NaCl, 10 mM imidazole, 5 mM 2-mercaptoethanol) supplemented with a cOmplete EDTA free protease inhibitor cocktail tablet (Roche) and lysed by sonication (MISONIX, 2 s on / 6 s off, 100 s total on time at 80% amplitude). The lysate was clarified by centrifugation at 41,656 × g for 30 min at 4 °C in a F21-8×50y rotor (Thermo Fisher Scientific). The clarified lysate was loaded to a 2.5 mL Ni-NTA (Qiagen) gravity flow column equilibrated with lysis buffer. The column was washed with 50 mL wash buffer 1 (50 mM HEPES-KOH pH 7.5, 1000 mM NaCl, 20 mM imidazole, 5 mM 2-mercaptoethanol) and 20 mL wash buffer 2 (50 mM HEPES-KOH pH 7.5, 300 mM NaCl, 5 mM 2-mercaptoethanol). Bound proteins were eluted with a buffer containing 50 mM HEPES-KOH pH 7.5, 300 mM NaCl, 200 mM imidazole and 5 mM 2-mercaptoethanol. The eluate was diluted with 4 volumes of buffer Q-A (50 mM Tris-HCl pH 8.0, 5 mM 2-mercaptoethanol) and loaded to a 5 mL HiTrap Q HP column equilibrated with buffer Q-AB (50 mM Tris-HCl pH 8.0, 100 mM NaCl, 5 mM 2-mercaptoethanol). The protein was eluted by applying an 80 mL linear gradient from 100% buffer Q-AB to 100% buffer Q-B (50 mM Tris-HCl pH 8.0, 500 mM NaCl, 5 mM 2-mercaptoethanol). The 6×His-MBP tag was cleaved with TEV protease and removed by flowing through the samples on a Ni-NTA column. The purified Ded1p was dialyzed twice against the storage buffer (10 mM HEPES-KOH pH 7.3, 200 mM KOAc, 2 mM DTT, 50% glycerol) and stored at -80ºC after liquid nitrogen freezing. The ATPase activity of the protein was verified by a malachite green assay as described (*49*).

**2. Yeast ribosomal subunits**

We have previously labeled and characterized the ybbR-tagged yeast 40S subunit (at the N-terminus of uS19) and 60S subunit (at the C-terminus of uL18) (*21*). Here, for double tagging and labeling of the 40S subunit, the S6 tag (amino acid sequence of GDSLSWLLRLLN) was genetically fused to the N-terminus of uS19 using the same strategy as described for the ybbR tagging of uS19. The resulting yeast strain was transformed with a dsDNA carrying a *URA3* cassette to tag the C-terminus of uS5 with the A1 tag (amino acid sequence of GDSLDMLEWSLM) via homologous recombination using standard PCR-based methods (*50*), and selected on SC-His-URA plates. Ribosome subunits were purified as previously described (*21*). A two-step labeling protocol was developed to obtain the Cy3.5-S6-uS19 and uS5-A1-Cy5 doubly labeled 40S subunits (*51*). First, 1 µM of the purified double-tagged 40S subunit was mixed with 2 µM SFP synthase and 10 µM Cy3.5-CoA in a reaction buffer containing 50 mM HEPES-KOH pH 7.5, 10 mM MgCl_2_ and 1 mM DTT. The reaction was incubated at 30ºC for 10 min before the sample was loaded on top of a sucrose cushion (containing 30 mM HEPES-KOH pH 7.5, 100 mM KOAc, 5 mM Mg(OAc)_2_, 2 mM DTT and 0.5 M sucrose). The ribosomes were pelleted by ultracentrifugation at 351,955 × g and 4 °C for 60 min in a TLA100.2 rotor, and the resulting Cy3.5-uS19-40S was resuspended in storage buffer (50 mM HEPES-KOH pH 7.5, 100 mM KOAc, 3 mM Mg(OAc)_2_, 1 mM DTT and 0.25 M sucrose). Next, 1 µM of Cy3.5-uS19-40S was mixed with 10 µM AcpS and 30 µM Cy5-CoA in the same reaction buffer as above, and the reaction mixture was incubated at 30ºC for 30 min. The ribosomes were pelleted by ultracentrifugation as described above and the final Cy3.5-Cy5-40S were resuspended in the storage buffer and stored at -80ºC after liquid nitrogen freezing. Site-specific labeling was verified by SDS-PAGE analysis of the samples from each labeling step (**fig. S3A**).

**3. Aminoacyl-tRNAs and mRNAs**

The native yeast Met-tRNA_i_ was from tRNA Probes, LLC (MI-60), and *E. coli* Phe-tRNA^Phe^ and Lys-tRNA^Lys^ labeled with Cy3.5, Cy5 or Cy5.5 were prepared and characterized as previously described (*49*). All the mRNAs used in this work were T7 RNA polymerase transcripts with DNA plasmids synthesized by GenScript USA Inc. or gblocks synthesized by Integrated DNA Technologies, Inc.. DNA templates were PCR-amplified from plasmids or gblocks, and used for in vitro transcription with MEGAscript T7 Transcription Kit (Invitrogen, #AM1334). The mRNA transcripts were purified using GeneJET RNA Purification Kit (Thermo Scientific, #K0731), and biotinylated at the 3′end and m^7^G-capped at the 5′ end by Vaccinia capping system (NEB, #M2080S) as previously described (*21*). The mRNA sequences were listed in **Table S1**.

When labeling the mRNAs by hybridization with a Cy3-DNA oligonucleotide (synthesized by Qiagen or Integrated DNA Technologies, Inc.), 200 nM mRNA and 100 nM DNA oligonucleotide were mixed in 50 mM Bis-Tris Propane pH 7.0 and 100 mM KOAc; the reaction mixture was heated at 65ºC for 2 min followed by cooling on ice. For each experiment, the mRNA–DNA hybrid was freshly prepared. To verify that the DNA was annealed to the expected position on the mRNA, the samples were treated with RNase H (M0297S, NEB) at 30ºC for 30 min and analyzed by 10% TBE-urea PAGE (see **fig. S5B** for an example).

For covalent labeling of mRNAs, a three-part splint ligation protocol was applied (*52*). The covalently labeled mRNAs differed in their 5′ parts (fragment 1) while shared the same two downstream fragments (fragments 2 and 3). The fragments 1-3 sequences for all the different ligated mRNAs were listed in **Table S1**. The T7 transcripts of the fragments 1 were m^7^G-capped by Vaccinia capping system, and sequentially treated with RNA 5′ Polyphosphatase (Lucigen #RP8092H) and Terminator 5′-Phosphate-Dependent Exonuclease (Lucigen #TER51020) to deplete uncapped transcripts. The T7 transcript of the fragment 3 was treated with RNA 5′ Polyphosphatase to obtain a mono-phosphate group at its 5′ end for subsequent ligation. The 3′end of the fragment 3 was biotinylated by periodate oxidation and hydrazide conjugation as described (*21*). To ligate the three parts of the mRNAs, 6 µM m^7^G-capped fragment 1, 2 µM fragment 2 (which contained a 5′ mono-phosphate group and an internal Cy3 dye), 4 µM biotinylated fragment 3 and 3 µM splint DNA (sequences listed in **Table S1**) were mixed in ddH_2_O. The sample was heated at 65ºC for 3 min, followed by 25ºC for 5 min and on ice for 5 min. Then the sample was mixed with 1 mM ATP and 1 U/µL T4 RNA ligase (Thermo Scientific, #EL0021) in a buffer containing 50 mM Tris-HCl pH 7.5, 10 mM MgCl_2_ and 10 mM DTT. The reaction mixture was incubated at 37ºC for 30 min, then Turbo DNase (Invitrogen #AM2238) was added to final 0.05 U/µL and the reaction was further incubated at 37ºC for 15 min to digest the splint DNA. The final mRNA was purified with MEGAclear Transcription Clean-Up Kit (Invitrogen #AM1908). **Fig. S3C** shows the *RPL30* mRNA ligation protocol workflow as an example.

**4. Single-molecule assays and data analyses**

All single-molecule experiments were performed on a customized ZMW-based RSII instrument from Pacific Bioscience (*24*). All the reaction mixtures for the single-molecule assays were prepared as outlined in **fig. S1A**, in a buffer containing 30 mM HEPES-KOH, pH 7.5, 100 mM KOAc and 3 mM Mg(OAc)_2_. A 3× eIF2•GTP•Met-tRNA_i_ ternary complex mixture was prepared by incubating 2.8 µM eIF2 with 1 mM GTP:Mg^2+^ at 30ºC for 10 min, then mixed with 2.7 µM Met-tRNA_i_ and incubated for another 5 min at 30ºC. Separately, the eEF1A•GTP•elongator tRNA ternary complexes were prepared by incubating 10 µM eEF1A, 6 µM eEF1Bα with 1 mM GTP:Mg^2+^ at room temperature (RT) for 5 min, followed by another 5 min incubation at RT after addition of 1.6 µM elongator tRNA (aminoacylated and with the appropriate dye label), resulting in an elongator TC at 1.6 µM (by tRNA). Next, the 43S PIC mixture was prepared by mixing 0.24 µM 40S (with the appropriate dye label), 3.6 µM eIF1, 3.6 µM eIF1A, 1× eIF2•GTP•Met-tRNA_i_ ternary complex, 0.6 µM eIF3, 1.8 µM Hcr1p, 1.2 µM eIF5 and 1 mM GTP:Mg^2+^, and incubated at 30ºC for 5 min.

For reaction Scheme 1 (**fig. S1A**), a delivery mixture was prepared by diluting the 43S PIC mixture (with Cy3-uS19 labeled 40S) by 3× with the addition of 0.1 µM Cy3.5-Phe-TC, 1.2 µM eIF4A, 1.2 µM eIF4B, 1.2 µM eIF4E, 1.2 µM eIF4G, 1.2 µM Ded1p, 1.2 µM Pab1p, 2 µM eIF5B, 0.2 µM Cy5-60S, 1 mM ATP:Mg^2+^ and 1 mM GTP:Mg^2+^, supplemented with 100 µg/mL casein (Sigma Aldrich, #C4032-100MG) as a blocking reagent to prevent non-specific binding of labeled components to the imaging surface, and an oxygen scavenging system (Pacific Bioscience) containing 2.5 mM of protocatechuic acid (PCA), 2.5 mM of TSY, and 2× PCD (protocatechuate-3,4-dioxygenase). Meanwhile, a ZMW chip (Pacific Bioscience) was treated and coated with Neutravidin as previously described (*21*). The unlabeled *RPL30* mRNA was immobilized on the imaging surface by incubating 30 µL mRNA at 0.2 nM in the ZMW chip at RT for 10 min. The chip was washed twice with an imaging buffer containing ATP:Mg^2+^, GTP:Mg^2+^, casein and the oxygen scavenging system at the same concentrations as in the delivery mix. To start the experiment, 20 µL of fresh imaging buffer was added to the ZMW chip, and the chip was loaded to the RSII instrument. Data acquisition at 10 frames per second and at 30ºC (or otherwise noted) were started with a single 532-nm laser excitation at 0.32 µWµm^-2^ and the delivery of 20 µL delivery mixture to the ZMW chip.

For reaction Scheme 2 (**fig. S1A**), the delivery mixture was the same as above but with eIF4A, eIF4E, eIF4G, Ded1p and Pab1p omitted, while the final imaging buffer was supplemented with eIF4A (at denoted concentrations), 1.2 µM eIF4B, 1.2 µM eIF4E, 1.2 µM eIF4G, 1.2 µM Ded1p and 1.2 µM Pab1p. From addition of the factor-containing imaging buffer to the chip to the beginning of the movie acquisition, the mRNA on the surface was preincubated with the factors for ~10 min.

For reaction Scheme 3 (**fig. S1A**), the imaging buffer was the same as that for Scheme 2 but without ATP, while the delivery mix contained 2 mM ATP, thus final ATP concentration was 1 mM after delivery mixture was added to the surface.

When titrating ATP concentrations or using slowly hydrolyzabe ATP analogs, the appropriate concentrations of the analogs were indicated in the main text and figures. Similarly, certain factors were omitted from the reactions as denoted in the text and figures to assess their effects on initiation.

For the four-color scanning assay (**Fig. 2, A to C**), reaction Scheme 2 was applied, but with the elongator TC omitted. The Cy3.5-uS19 and uS5-Cy5 doubly labeled 40S and uL18-Cy5.5-60S were used. The mRNAs were Cy3-labeled as described above. When using mRNAs labeled by Cy3-oligonucleotide hybridization, Ded1p was omitted from the imaging buffer. This was because that the oligonucleotide would be rapidly displaced by Ded1p in the presence of ATP before the movie acquisition and the start of the reaction (*19*), leading to loss of the mRNA signal. The two-color scanning assay (**Fig. 3**, **A and B**) was performed in the same way as the four-color scanning assay, except that the uS5-Cy5-40S and unlabeled 60S were used. To determine the mRNA-uS5 FRET efficiencies with varying 40S to mRNA-Cy3 label distances (**fig. S3B**), the two-color scanning assay setup was used, with the *RPL30* mRNA labeled by Cy3-DNA oligonucleotides hybridized to the designated positions on the mRNA. To measure upstream CUG initiation efficiency in relation to that of the AUG initiation (**Fig. 5B**), reaction Scheme 2 was followed. Here, unlabeled mRNAs, unlabeled 40S and uL18-Cy3-60S were used, and the delivery mixture contained both Cy5.5-Phe-TC and Cy5-Lys-TC.

The acquired four-color movies were analyzed using custom MATLAB (Mathworks) scripts as described (*21*). Briefly, the fluorescence traces from individual ZMWs were manually examined for the presence of the single fluorophores of interests at different time points (e.g. the signal from immobilized, labeled mRNA was expected to be present at the beginning of the movie, whereas signals from fluorophores attached to the 40S, 60S or elongator tRNAs were expected to appear later in the movie). The individual association and dissociation events were manually assigned according to the appearance and disappearance of the corresponding fluorescence signals, to obtain the duration of each molecular event. The mean lifetime (± 95% confidence interval) of each molecular event was estimated by fitting the cumulative probability distribution to a single- or double-exponential equation using cftool in MATLAB. The relationship between the mean scanning time and the 5′UTR length was fit to a linear equation (**Fig. 2E**) using Prism version 6.0 (GraphPad Software Inc.). Two-tailed unpaired Student’s *t*-test was performed using Prism for the scanning times determined under varying ATP conditions with *RPL30* and *PMA1* mRNAs (**Fig. 3I**).

For data from the two-color scanning assay, the Cy3 and Cy5 fluorescence traces were corrected for fluorescence background and bleed-through on a trace-by-trace basis, and the FRET vs. time trajectories were obtained by plotting the time-evolution of the FRET efficiency (*E*_FRET_ = *I*_Cy5_/(*I*_Cy5_ + *I*_Cy3_), where *I*_Cy3_ and *I*_Cy5_ are the intensities of the donor and acceptor dye intensities). The FRET trajectories were idealized with a two-state hidden Markov model implemented with ebFRET followed by manual inspection to correct mis-assigned events (the FRET ramp was treated as the low FRET state) (*53*). Distribution histograms of the *E*_FRET_ values were plotted in MATLAB and fit to single or double gaussian model to obtain the mean *E*_FRET_ (± standard deviation) of the FRET states. To estimate the mean lifetime of the FRET ramp, all the FRET trajectories from each experiment were post-synchronized with the beginning of the FRET set to time 0 and averaged at each movie frame; then the averaged *E*_FRET_ vs. time trajectory, which mimics the ensemble averaged behavior of all the analyzed molecules, was fit to a single-exponential equation as previously described (*54*). To determine the kinetics of the FRET state transitions in experiments with mRNAs carrying 5′UTR structures of different stabilities, the cumulative probability distributions of the state durations were fit to a single- or double-exponential equation, and the estimated mean lifetimes were reported with the 95% confidence intervals.

**5. Structural models**

Ribosome models (**Fig. 2B** and **fig. S11**) were analyzed and visualized by ChimeraX (*55*) with PDB 6TNU (*56*).

**6. Re-analysis of the human 40S profiling data**

To determine the occupancy of 40S ribosomes on and around structures in proximity to the start codon of endogenous human mRNAs, sequences at positions -55 to -5 relative to the start codon were extracted from Ensembl transcript assembly 94 for 5′UTRs longer than 55 nucleotides. The minimum free energies of these 50-mers were calculated using RNAfold (*57*). The transcripts were grouped into two buckets: bucket 1 (referred to as “stable” in **fig. S13**) with the 50-mers having the potential to form secondary structures more stable than -20 kcal/mol (n = 4890) and bucket 2 (referred to as “weak” in **fig. S13**) with those less stable (n = 28217). Then, the 40S ribosome footprinting data from Bohlen *et al*. was re-analyzed to determine the frequency of 40S ribosomes on and around this structured/unstructured area using transcriptome alignments as described (*44*), which were further processed with custom C++ scripts available at Teleman AG Github (<https://github.com/aurelioteleman/Teleman-Lab>). Transcripts with PCR artefacts within this area of the mRNA were excluded from the analysis, leaving 4677 transcripts in bucket 1 and 27765 transcripts in bucket 2. Finally, 40S footprint counts were normalized to the sequencing depth (number of reads aligned to the transcriptome) for each independent library and compared.

**Supplementary Figures and Legends:**

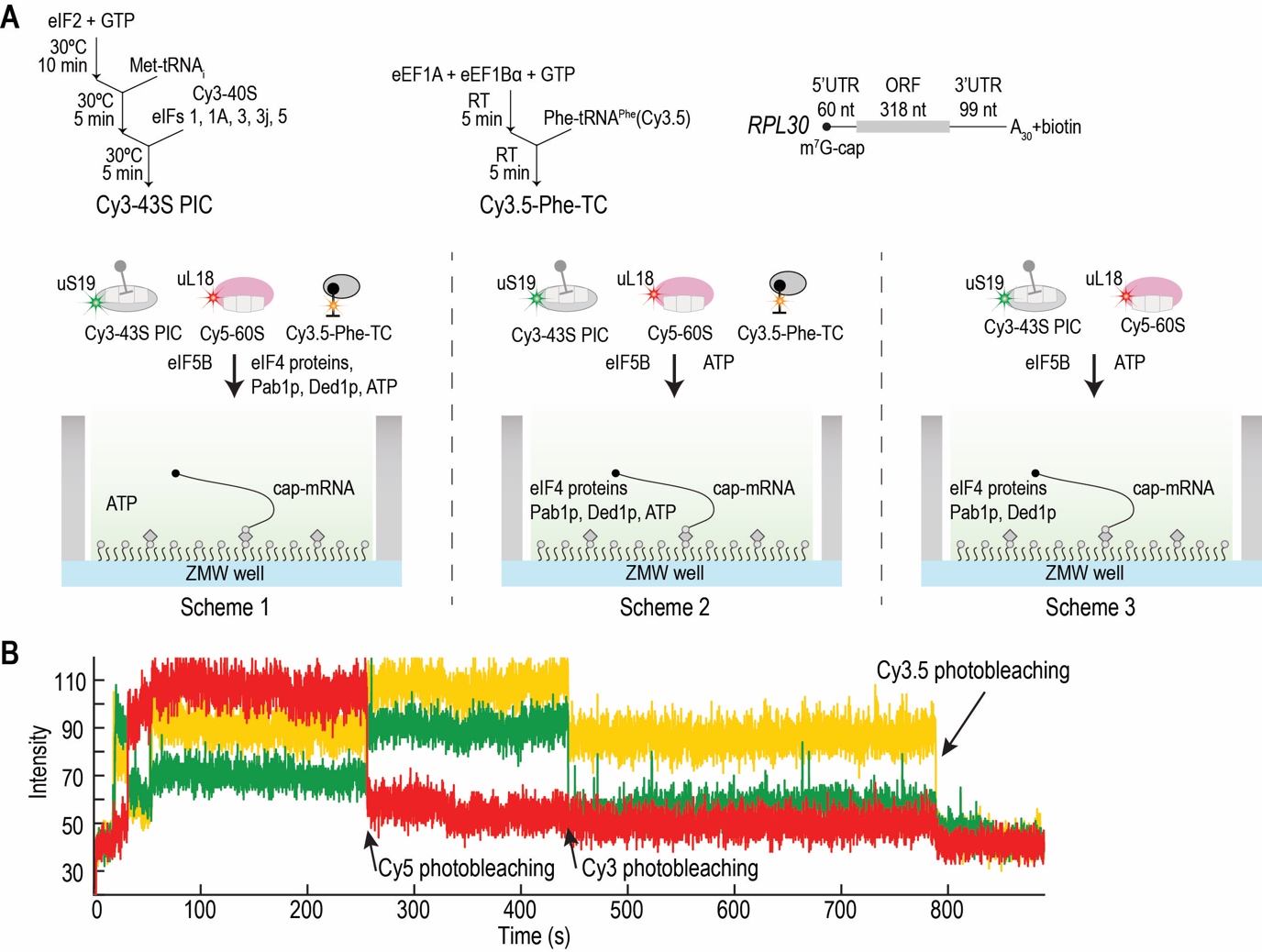

**Fig. S1. Direct observation of initiation in real time.**

(**A**) Reaction scheme and setup. Top, the workflows to pre-form a 43S PIC mixture and an elongator tRNA ternary complex (TC) mixture, and a cartoon illustration of the full-length, 5′ m^7^G-capped, 3′ biotinylated *RPL30* mRNA (ORF, open reading frame). Bottom, three different reaction schemes to track initiation. Scheme 1, all the other components (including 1 mM ATP) were co-delivered to surface-tethered single mRNAs upon laser excitation and start of movie acquisition (1 mM ATP was also included in the imaging buffer on the surface); Scheme 2, eIF4 proteins (including eIF4A, eIF4G, eIF4E and eIF4B), Pab1p, Ded1p and ATP were included in the imaging buffer and pre-incubated with the tethered mRNAs for ~10 min before the delivery of all the other components; Scheme 3, similar to Scheme 2, but with ATP only present in the delivery mixture and Phe-TC omitted. Final concentrations after mixing were: 600 nM eIF4A, 600 nM eIF4B, 600 nM eIF4E, 600 nM eIF4G, 600 nM Ded1p, 600 nM Pab1p, 40 nM Cy3-40S PIC (by 40S, with 600 nM eIF1, 600 nM eIF1A, 150 nM eIF2•GTP•Met-tRNA^i^ ternary complex, 100 nM eIF3, 300 nM eIF3j, 200 nM eIF5), 1 µM eIF5B, 100 nM Cy5-60S, 50 nM Cy3.5-Phe-TC (by Cy3.5-Phe-tRNA^Phe^), 1 mM ATP:Mg^2+^ and 1 mM GTP:Mg^2+^, unless otherwise stated. (**B**) The full example fluorescence vs. time trace for **Fig. 1D**.

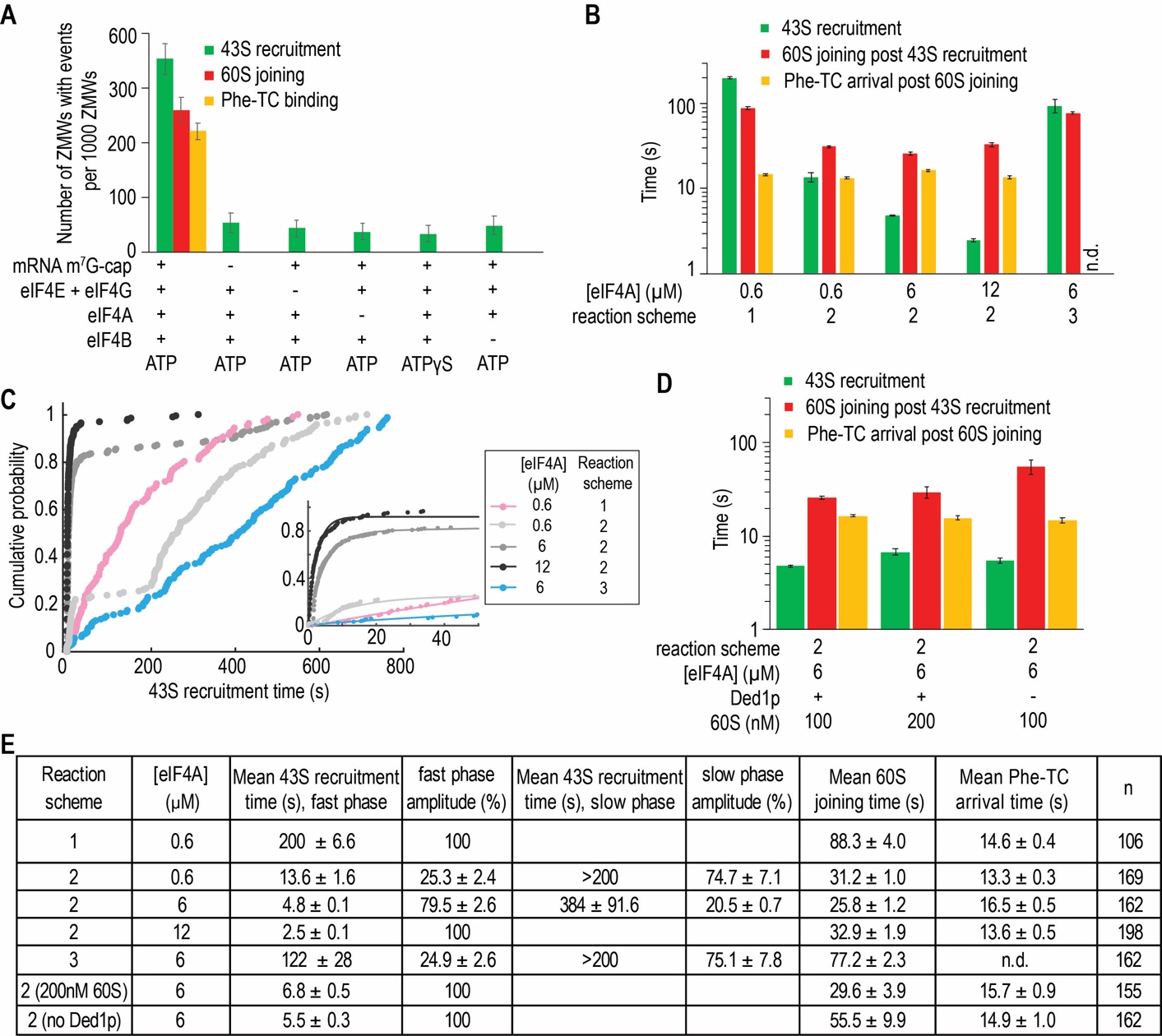

**Fig. S2. High eIF4A concentration and ATP hydrolysis drive rapid 43S PIC recruitment.**

(**A**) With reaction Scheme 1 (**fig. S1A**) and at 0.6 μM eIF4A concentration, 35.3 ± 2.9% (mean ± 95% CI) of the ZMWs contained one 40S binding event to *RPL30* mRNA (in the range that maximize the likelihood of single complex loading per ZMW well according to Poisson statistics, n = 1000), 73.6 ± 6.2% led to 60S joining to form the 80S, and 85 ± 5.7% the 80S complex accepted the first elongator Phe-TC and thus progressed to the elongation phase. The 40S–mRNA binding was inhibited on the same mRNA lacking an m^7^G-cap, or when eIF4E and eIF4G, eIF4A, or eIF4B were omitted, or when 1 mM ATPγS was used instead of ATP (only 5.3 ± 1.8%, 4.3 ± 1.5%, 3.7 ± 1.5%, 4.8 ± 1.7%, 3.3 ± 1.5% of ZMWs had one 40S event, respectively, and none of these events progressed to 60S joining; n = 1000). These assured that the reactions we observed arose mainly from the canonical cap-dependent pathways. (**B-E**) The kinetics of individual initiation steps under different reaction conditions or schemes. Kinetic curves were fit to single- or double-exponential equations (only the fast phase mean times used for bar plots). Data were shown as mean ± 95% CI, and n.d. means not determined. [**Note**: With reaction Scheme 1 and eIF4A at 0.6 μM, the mean time of 43S recruitment was ~200 s (at 40 nM Cy3-43S PIC concentration), the subsequent 60S joining ~88 s (at 100 nM Cy5-60S concentration), and the transition to elongation (i.e. Phe-TC arrival) ~15 s (at 50 nM Cy3.5-Phe-TC) (**B**, **C**, **E**). With reaction Scheme 2, the surface-tethered mRNAs were pre-incubated with eIF4 proteins, Ded1p, Pab1p and ATP for ~10 min before the start of the reaction. At 0.6 µM eIF4A, this pre-incubation resulted in ~25% of the total 43S recruitment events occurring at a fast timescale (~13.6 s) while the other ~75% at ~200 s. Raising eIF4A concentrations to 6 and 12 µM increased both the proportion and rate of fast 43S recruitment events (~80% and 100%, ~4.8 s and 2.5 s, respectively). The stimulatory effect required the presence of ATP during the pre-incubation with the mRNA, as reaction Scheme 3 led to slow 43S PIC recruitment with ATP only delivered after the pre-incubation. Thus, all our other experiments were performed with the mRNAs pre-incubated with eIF4 proteins and ATP, unless otherwise stated.]

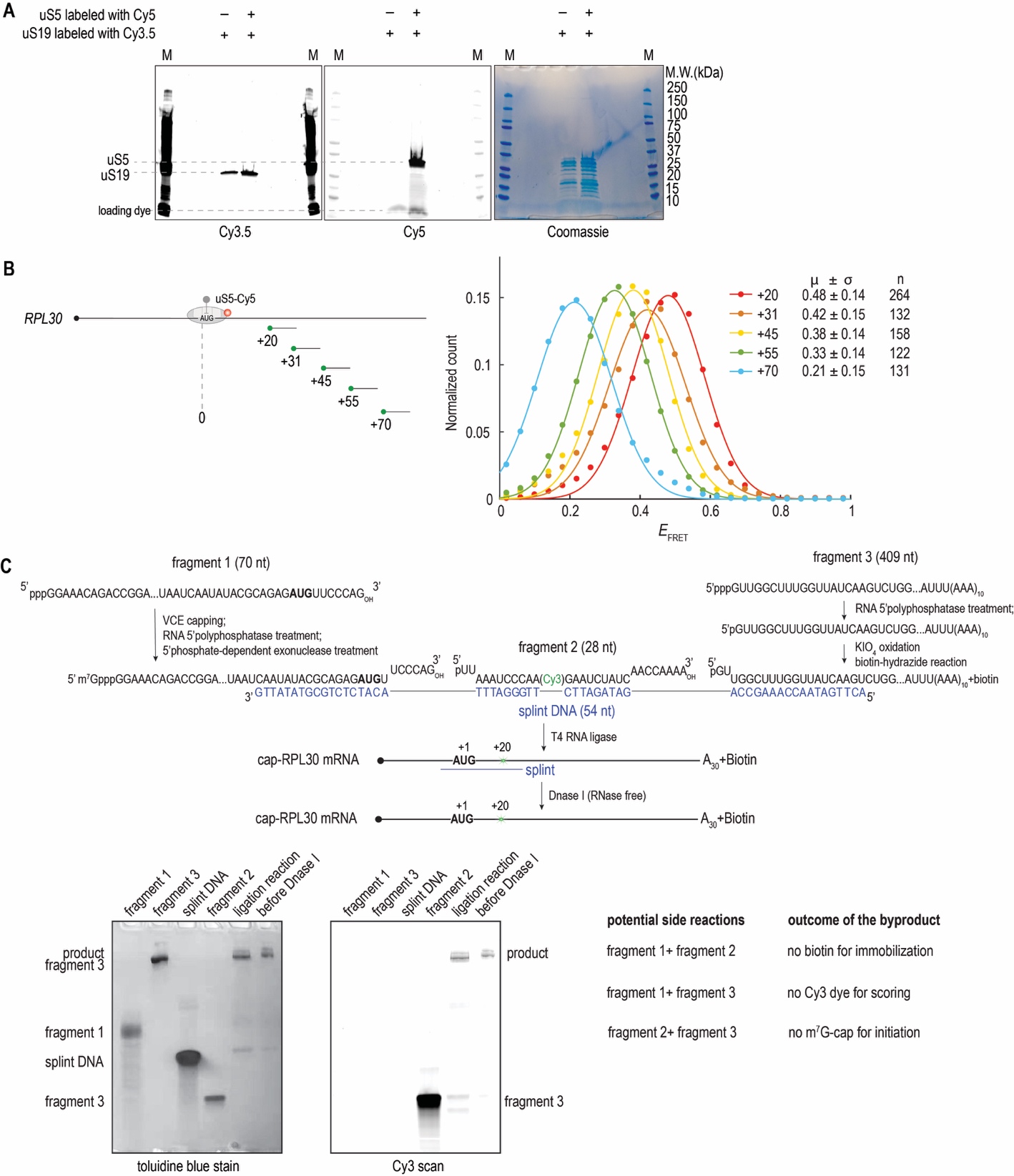

**Fig. S3. Labeling of ribosomes and mRNAs for direct observation of scanning.**

(**A**) Example SDS-PAGE (4-20%) analysis of dye labeled 40S subunits. The 40S was first labeled with Cy3.5-CoA at the S6-tag fused to N-terminus of uS19 by SFP synthase, purified, and then labeled with Cy5-CoA at the A1-tag fused to the C-terminus of uS5 by AcpS. The fluorescent signals from the bands detected confirmed site-specific dual labeling of the 40S subunits. (**B**) Calibration of the Cy3-mRNA to uS5-Cy5 FRET efficiency (*E*_FRET_) with varying ribosome to Cy3-oligonucleotide distances. The *RPL30* mRNA was labeled at different positions downstream of the AUG codon via hybridization with Cy3-DNA oligonucleotides. Using reaction Scheme 2 (**fig. S1A**) with Ded1p omitted (as Ded1p would rapidly remove the DNA oligonucleotide from the mRNA) and with uS5-Cy5-40S, the *E*_FRET_ between mRNA and uS5 were measured and the distribution histograms were fit to a single Gaussian equation to obtain the mean *E*_FRET_ (μ) ± standard deviation (σ) values. As expected, *E*_FRET_ between uS5-Cy5 and Cy3-mRNA was dependent on the distance between the ribosome position to the Cy3-dye position on the mRNA. (**C**) A schematic workflow and design for three-part splint ligation to introduce a site-specific, covalent Cy3-label at the +20 position (with the A of AUG set to 0) in *RPL30* mRNA. The gel pictures show the analysis of the reaction intermediates by 10% TBE-urea PAGE. The potential byproducts from side reactions were also listed with the reasons why they should not affect the singe-molecule assays.

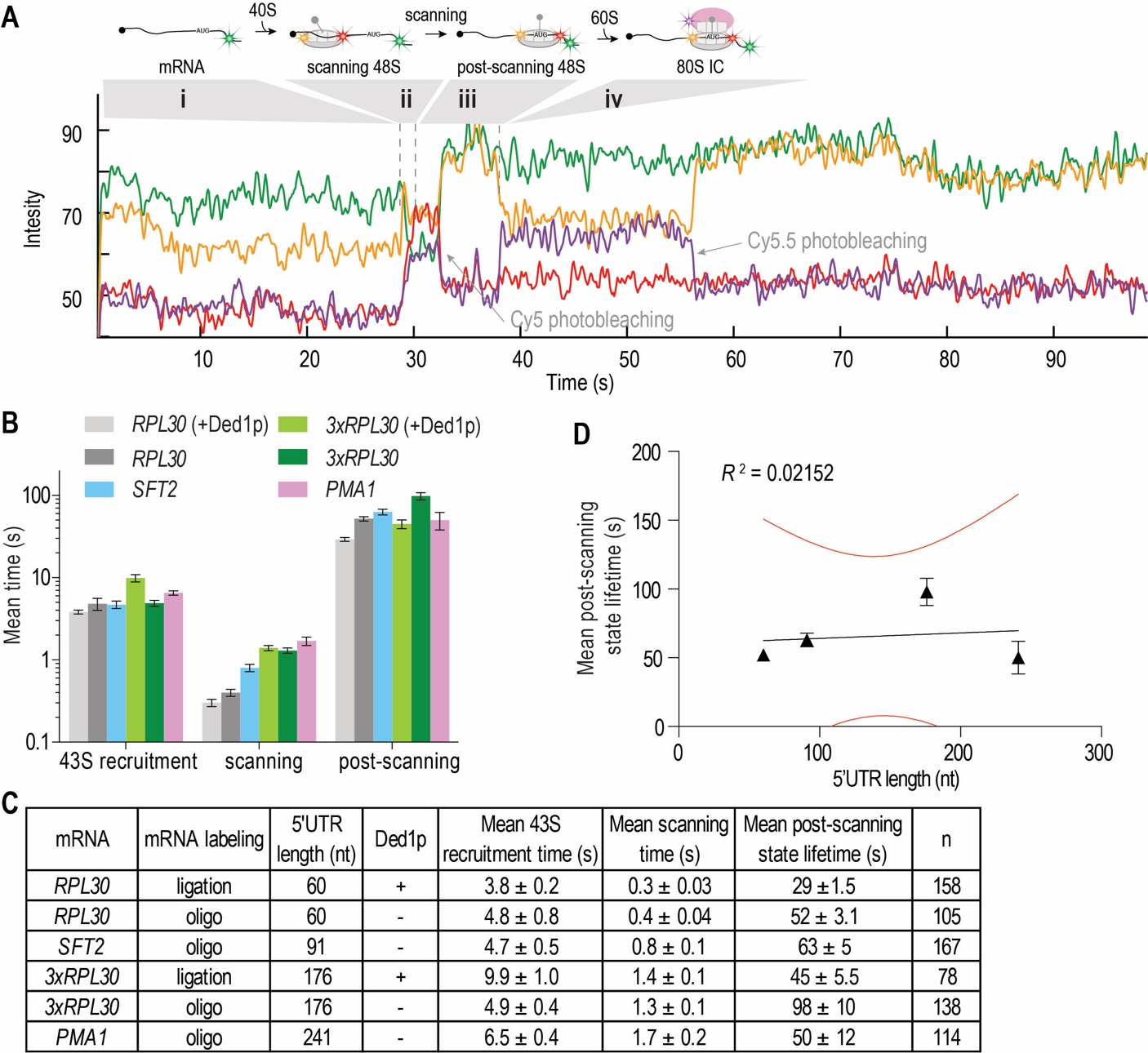

**Fig. S4. Observing scanning by two orthogonal FRET pairs.**

(**A**) The full trace for **Fig. 2C**, which demonstrated the dissection of initiation with high kinetic resolution: surface-tethered Cy3-mRNA was identified as the green signal (state **i**), recruitment of 43S PIC (also the start of scanning) was marked by the burst of the Cy3.5-uS19 signal (yellow, direct illumination by the green laser; state **i** to **ii** transition), end of scanning was shown by the Cy3-mRNA to Cy5-uS5 FRET reaching its maximum efficiency (beginning of state **iii**), and post scanning, the joining of Cy5.5-60S (purple, by FRET with Cy3.5-uS19; state **iii** to **iv** transition) concluded the initiation process. (**B** and **C**) The mean times for 43S PIC recruitment (state **i** lifetime), scanning (state **ii** lifetime), and the post-scanning state prior to 60S joining (state **iii** lifetime) from experiments performed with varying mRNAs in the presence (mRNA covalently labeled) or absence (mRNA labeled with Cy3-oligonucleotides) of Ded1p. (**D**) The mean post-scanning state lifetimes from experiments in the absence of Ded1p were plotted against the 5′UTR lengths and fit to a linear equation (black line, with red lines as 95% CI), which resulted in an *R*^2^ = 0.02152, indicating a non-linear relationship. Errors represent the 95% CI of the mean.

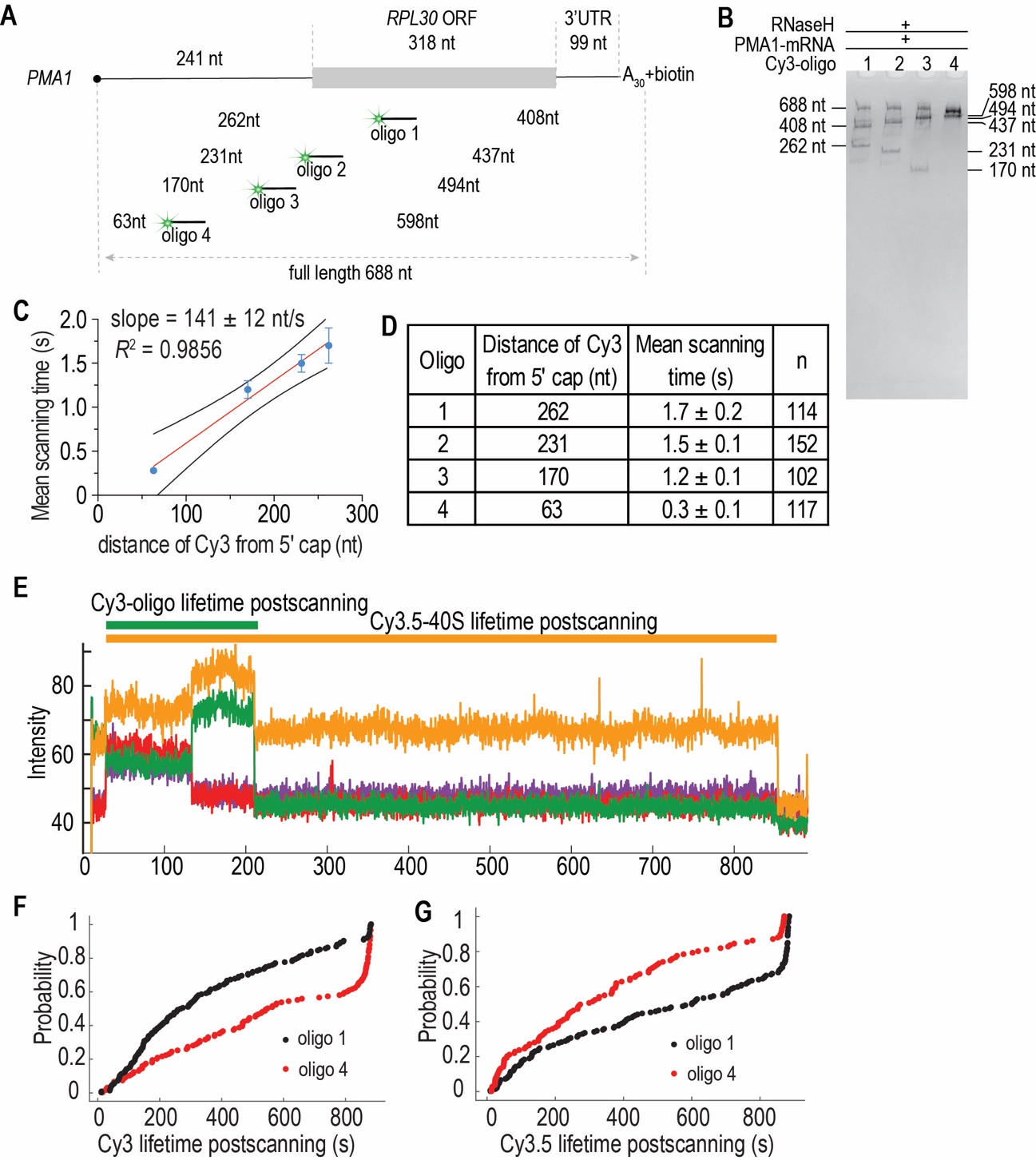

**Fig. S5. Scanning time increases linearly with the number of nucleotides to traverse in the 5′UTR.** (**A**) A series of Cy3-DNA oligonucleotides were hybridized separately to the *PMA1* mRNA at varying positions, leading to different distances from the 5′ end. The Cy3-oligo 1 was annealed to the +20 position, while the others to the 5′ UTR. (**B)** The site-specific annealing of the oligonucleotides was verified by RNase H treatment and analyzed on 10% TBE-urea PAGE. First, 200 nM *PMA1* was annealed with 100 nM Cy3-oligonucleotide in 50 mM Bis-Tris propane pH 7.0 and 100 mM KOAc (total volume of 10 μL) at 65ºC for 2 min and cooled on ice (same procedure followed when preparing labeled mRNAs for single-molecule assays). Then 0.5 μL of 5 U/μL RNase H was added to the reaction mixture followed by another 30 min incubation at 30ºC. The samples were analyzed on 10% TBE-urea PAGE and the pattern of the bands suggested specific annealing of all the oligonucleotides. (**C** and **D**) With *PMA1* labeled at different positions with the DNA oligonucleotides, the mean scanning times were determined in the four-color scanning assay (**Fig. 2A**) and plotted against the distance between the Cy3-dye and the 5′ end of the mRNA. Data were shown as mean ± 95% CI, the red line shows the linear fitting of the mean scanning time against Cy3–5′ end distance, with the 95% CI shown as the black curves. (**E**) Sample trace from experiment with Cy3-oligo 4 labeled *PMA1* mRNA. The Cy3.5-uS19 (direct illumination) and uS5-Cy5 (FRET with Cy3-oligo) signals from the double-labeled 40S subunit were observed, but not the Cy5.5-60S subunit signal. The cumulative probability distributions of the observed Cy3 lifetime (**F**) and Cy3.5 lifetime (**G**) after Cy3–Cy5 FRET efficiency reaching its maximal value were plotted for experiments with *PMA1* mRNA labeled with Cy3-oligo 1 (n = 241) and Cy3-oligo 4 (n = 115). [**Note**: Cy3-oligo 1 was not expected to be displaced from the mRNA by the initiating ribosomes, since it was located at position +20. Thus, the Cy3 and Cy3.5 lifetimes from experiments with Cy3-oligo 1 were used as positive controls showing stable DNA–mRNA and 40S–mRNA interactions. Cy3-oligo 4 was annealed to the 5′UTR, and similar Cy3 lifetime was observed as for Cy3-oligo 1, indicating that the anti-sense DNA targeting the 5’UTR was not dislodged during scanning. Meanwhile, the Cy3.5 lifetimes were also similar when the two DNA oligonucleotides were used, while 60S joining was not observed with Cy3-oligo 4. These suggested that the scanning 43S PIC was stalled by the Cy3-oligo 4, and remained associated with the mRNA.]

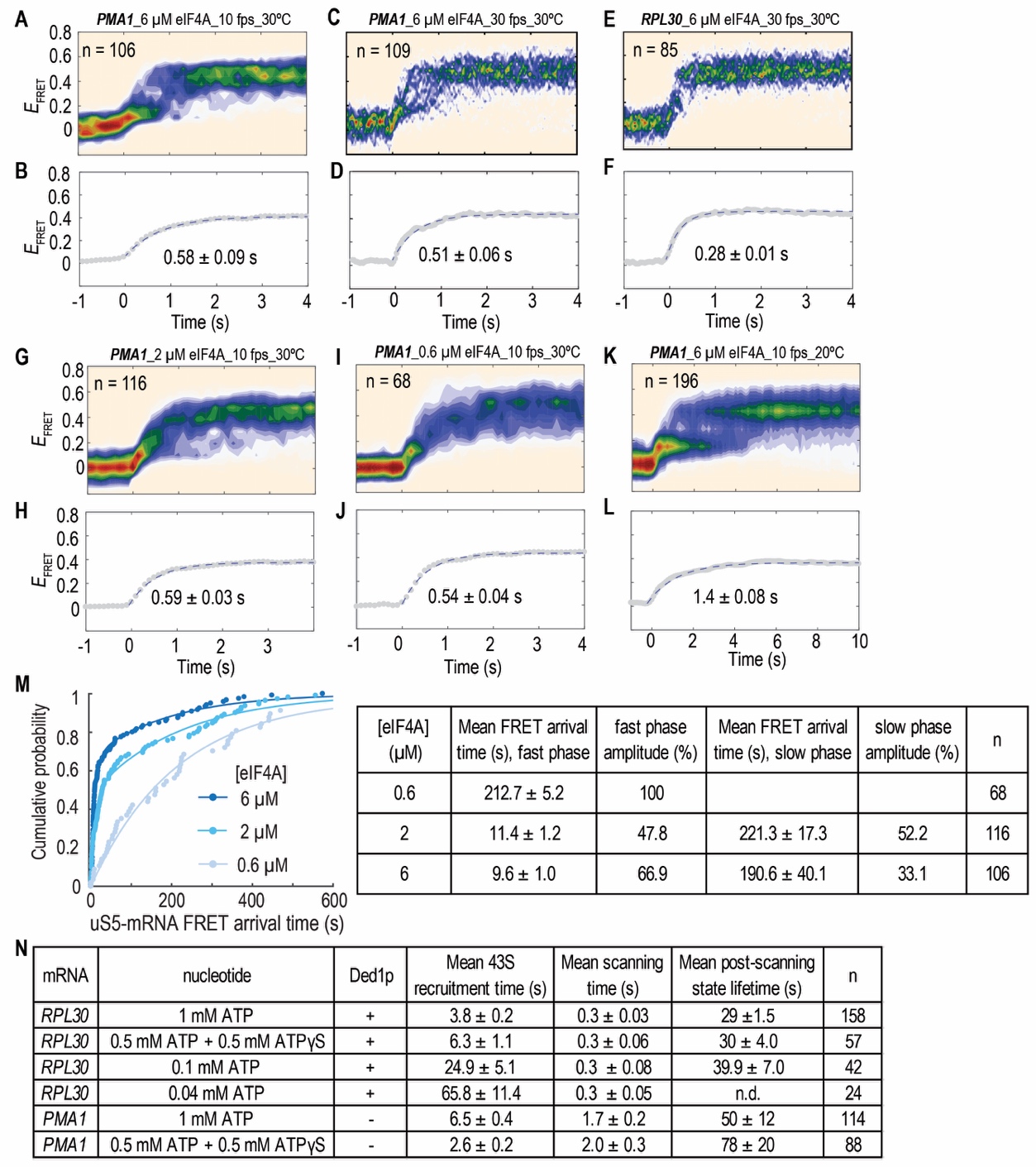

**Fig. S6. Scanning is independent of multiple rounds of ATP hydrolysis by eIF4A.**

(**A-L**) For two-color mRNA–uS5 FRET experiments with *PMA1* or *RPL30* mRNA at different eIF4A concentrations, reaction temperatures and movie frame rates, the *E*_FRET_ vs. time trajectories were post-synchronized with the beginning of the FRET ramp set at time 0 and plotted as density heat maps. The averaged trajectories were fit to a single-exponential equation to estimate the mean lifetimes of the FRET ramp (data shown as mean ± 95% CI). (**M**) Cumulative probability distributions of the observed mRNA–uS5 (Cy3–Cy5) FRET arrival times in the two-color FRET assays with *PMA1* mRNA at different eIF4A concentrations. The distributions were fit to a single- (for 0.6 μM eIF4A) or double- (for 2 and 6 μM eIF4A) exponential equation to estimate the mean arrival times. (**N**) Kinetics from the four-color scanning assays with *RPL30* and *PMA1* mRNAs under varying reaction conditions (n.d., not determined due to too few molecules for robust kinetics analysis). Errors represent the 95% CI of the mean.

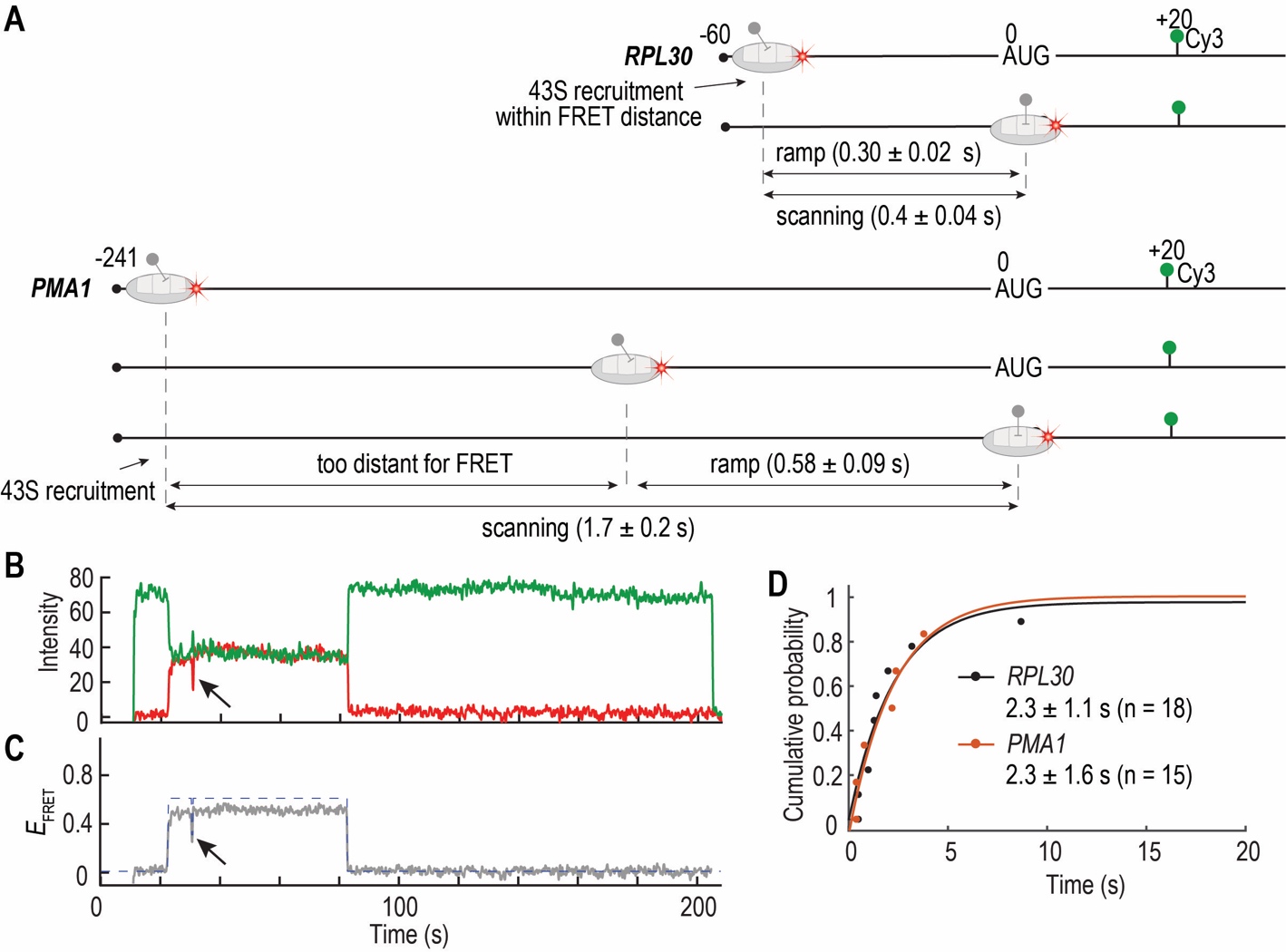

**Fig. S7. The 43S PIC landing site is near the 5′ region.**

(**A**) The 5′UTR of *RPL30* mRNA is 60 nt, and when the 43S PIC is recruited to the mRNA, the landing site is already within the FRET distance to allow mRNA–uS5 FRET. Thus, the FRET ramp lifetime was similar to the total scanning time determined with the four-color scanning assay. By contrast, the PMA1 mRNA has a 5′UTR of 241 nt, and the 43S PIC landing site is likely too distant for mRNA–uS5 FRET. Thus, there is a delay from 43S recruitment to when the mRNA–uS5 FRET starts to appear, and the FRET ramp lifetime was only a fraction of the total scanning time. (**B**) Sample fluorescence trace and (**C**) *E*_FRET_ vs. time trajectory showing the occasional backward (3′ to 5′) movement by the 43S PIC, as shown by the transition from the high mRNA–uS5 FRET state to a low state (indicated with black arrows). For *RPL30*, 10.8 ± 4.8% of the traces showed at least one FRET state transition (n = 18 out of 167), with average 0.14 ± 0.04 transition cycles per trace. For *PMA1*, 14.3 ± 7.6% of the traces showed at least one FRET state transition (n = 15 out of 105), with average 0.15 ± 0.04 transition cycles per trace. (**D**) The cumulative probability distribution of the observed lifetime of the low FRET state within fluctuations in experiments with *RPL30* and *PMA1* mRNAs. Data were fit to a single-exponential equation and data were shown as mean lifetime ± 95% CI.

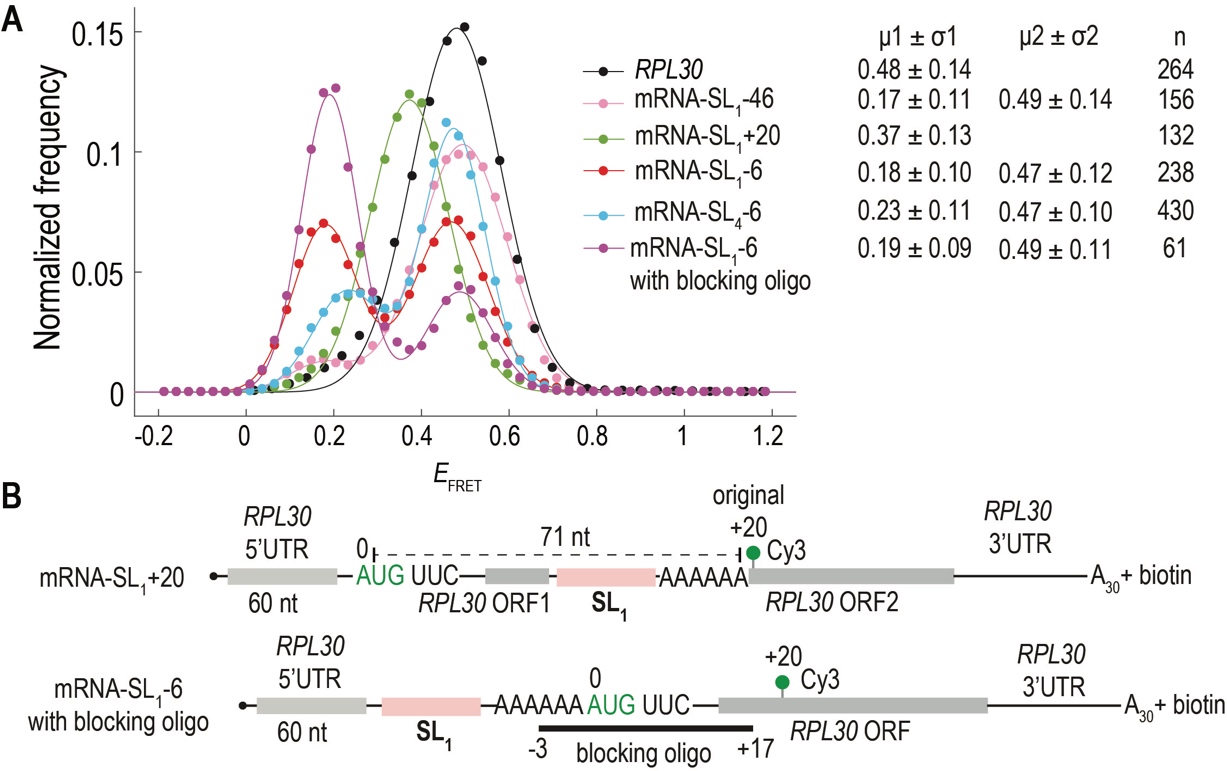

**Fig. S8. The high FRET state in mRNA–uS5 two-color FRET correlated with the 43S at the start site.** (**A**) The *E*_FRET_ distribution histograms from experiments with different mRNAs were fit to a single or double Gaussian equation, and the mean *E*_FRET_ (μ) ± standard deviation (σ) values were shown. [**Note**: With a mRNA bearing SL_1_ in between the AUG and the original +20 position (mRNA-SL_1_+20, also see panel **B** here), a lower *E*_FRET_ than that of *RPL30* was observed, demonstrating extra FRET distance contributed by the SL_1_ hairpin structure. The high FRET state efficiency observed for mRNA-SL_1_-46 was similar to that for *RPL30*, suggesting that the 40S reached the AUG site. Thus, whereas scanning is inhibited by anti-sense DNA oligonucleotides targeting the 5’UTR (**fig. S5, E-G**), SL_1_ located 46 nt upstream from the AUG did not significantly stall the scanning ribosome.] (**B**) Cartoon illustrations of the mRNA variants. Top, SL_1_ and the AAAAAA sequences were moved to the downstream of the start site at the original +20 position, thus splitting the *RPL30* ORF into two portions (mRNA-SL_1_+20). Bottom, an unlabeled blocking DNA oligonucleotide (Kozak-RPL30-AUG blocking oligo in **Table S1**) was annealed to the -3 to +17 position on the mRNA-SL_1_-6, along with the Cy3-oligonucleotide at +20 position.

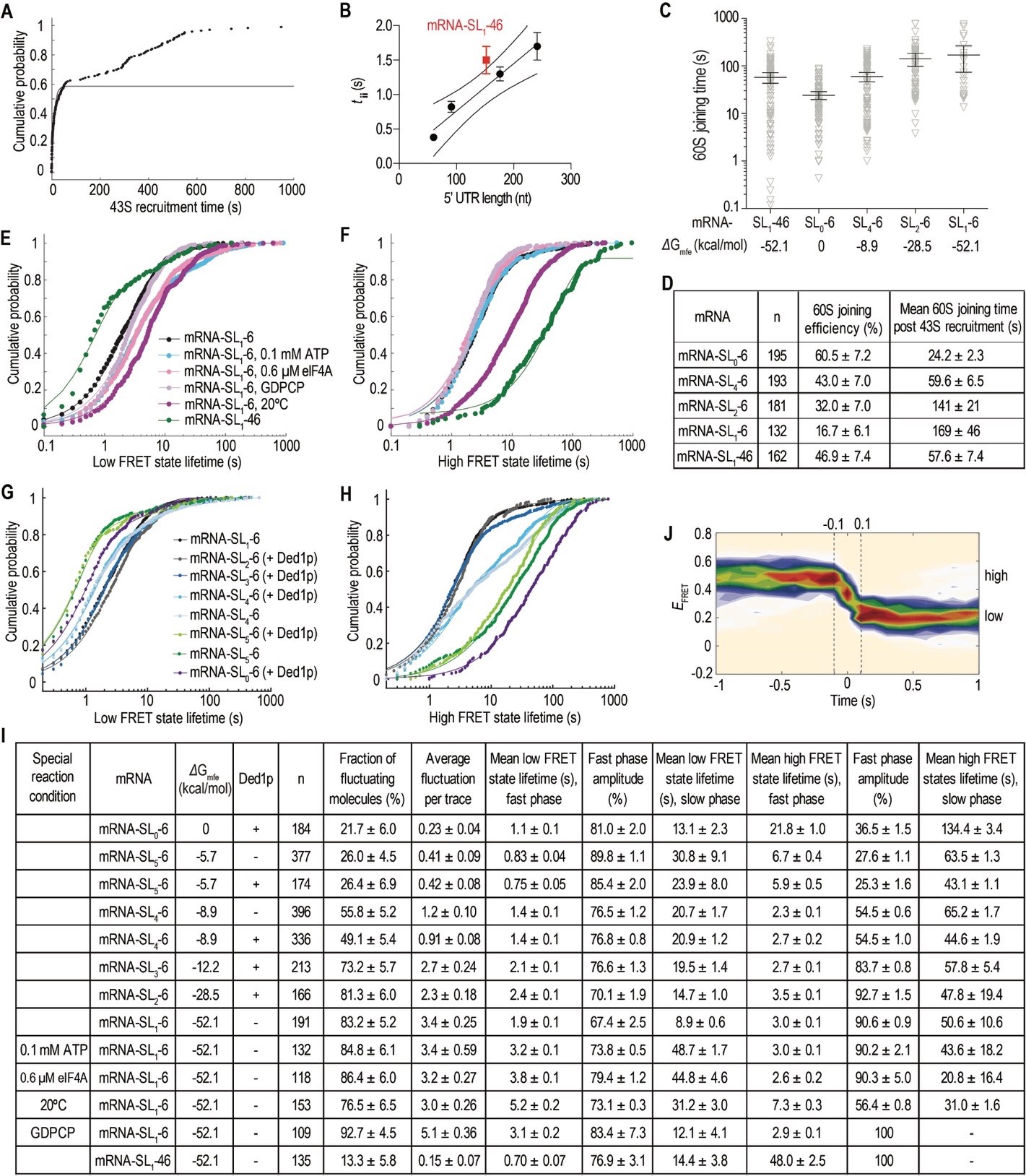

**Fig. S9. Kinetics of scanning and 60S joining efficiencies.**

(**A**) The cumulative probability distribution of the observed 43S recruitment time with mRNA-SL_1_-46 displayed two phases. The dominant fast phase accounted for 58.6 ± 1.7% amplitude with a mean time of 9.2 ± 0.9 s (mean ± 95% CI, n = 162). (**B**) The mean scanning time determined for mRNA-SL_1_-46 (red symbol and error bars represent mean ± 95% CI) was plotted together with **Fig. 2E** for comparisons. Here, the scanning speed was slightly smaller with mRNA-SL_1_-46 than the other mRNAs without stable structures in the 5′UTR. (**C**) The observed 60S joining times post 43S recruitment (open triangles) measured for mRNAs with hairpin structures of varying stabilities in the four-color scanning assay (**Fig. 2A**, in the absence of Ded1p). Error bars represent the 95% CI of the mean 60S joining times. (**D**) 60S joining efficiencies (calculated as the fraction of 48S PICs that progressed to 60S joining) and times determined from experiments as in (**C**). Data were shown as mean ± 95% CI. (**E**-**I**) Kinetic analysis of the mRNA–uS5 FRET fluctuations observed with different mRNAs under indicated conditions. Here, the mRNA-SL_1_-6 and mRNA-SL_1_-46 were labeled by hybridization with a Cy3-DNA oligonucleotide at +20 position, while the other mRNAs were covalently labeled via the ligation approach (see Methods). The canonical condition was at 6 μM eIF4A, 1 mM ATP, 1 mM GTP, 30ºC. The cumulative probability distributions of the observed low FRET state lifetimes (**E** and **G**) or high FRET state lifetimes (**F** and **H**) were fit to a single- or double-exponential equation to estimate the mean lifetimes. Data in (**I**) were mean ± 95% CI. (**J**) The mRNA–uS5 FRET fluctuation events from experiments with mRNA-SL_1_-6 were post-synchronized with the high to low state transition set at time 0. Dashed vertical lines marked the timepoints -0.1 and 0.1 s, which corresponded to the movie frames before and after the transition. Results indicated that the transition occurred within a single movie frame.

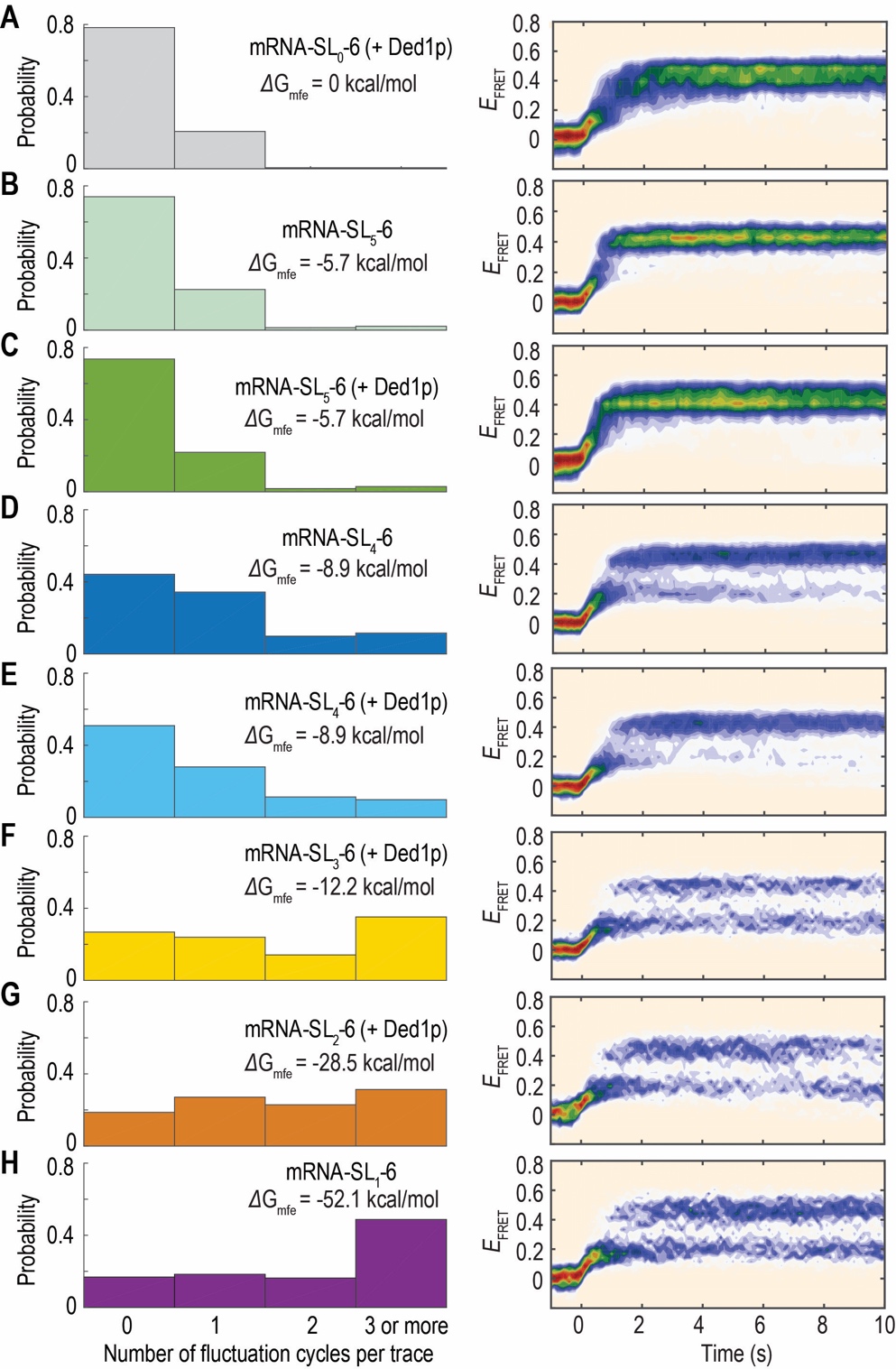

**Fig. S10. Stable hairpins in the 5′UTR induce mRNA–uS5 FRET fluctuations.**

(**A**-**H**) Left panels, the distributions of the number of FRET fluctuation cycles observed per *E*_FRET_ trajectory, with 3 and more cycles binned together. Right panels, all the FRET events were post-synchronized with the beginning of the events set at time 0. Same as in **fig. S9I**, the mRNA-SL_1_-6 and mRNA-SL_1_-46 were labeled by hybridization with a Cy3-DNA oligonucleotide at +20 position, while the other mRNAs were covalently labeled. Number of molecules analyzed were the same as in **fig. S9I**. Results demonstrated that the FRET fluctuations were more frequent with more stable hairpins in the 5′UTR, and thus with the low FRET state more populated in the post-synchronized heat map.

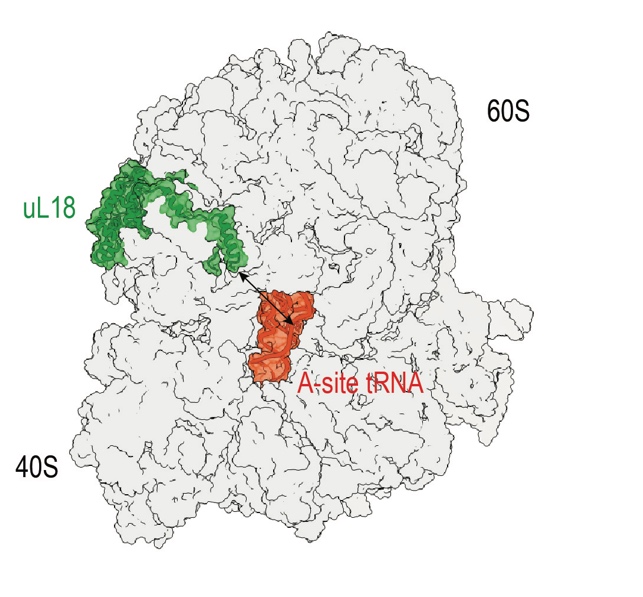

**Fig. S11. Distance between 60S subunit and A-site tRNA labeling sites.**

The 60S ribosomal subunit was labeled with a Cy3 donor dye at the C-terminus of uL18 (green), and the *E. coli* elongator tRNA^Phe^ and tRNA^Lys^ were labeled with Cy5.5 and Cy5 acceptor dyes, respectively, at the acp^3^U47 position. The distance between the labeling sites of the 60S subunit and the A-site tRNA (indicated by the black arrow) is predicted to be < 70 Å (PDB 6TNU), thus within FRET distance.

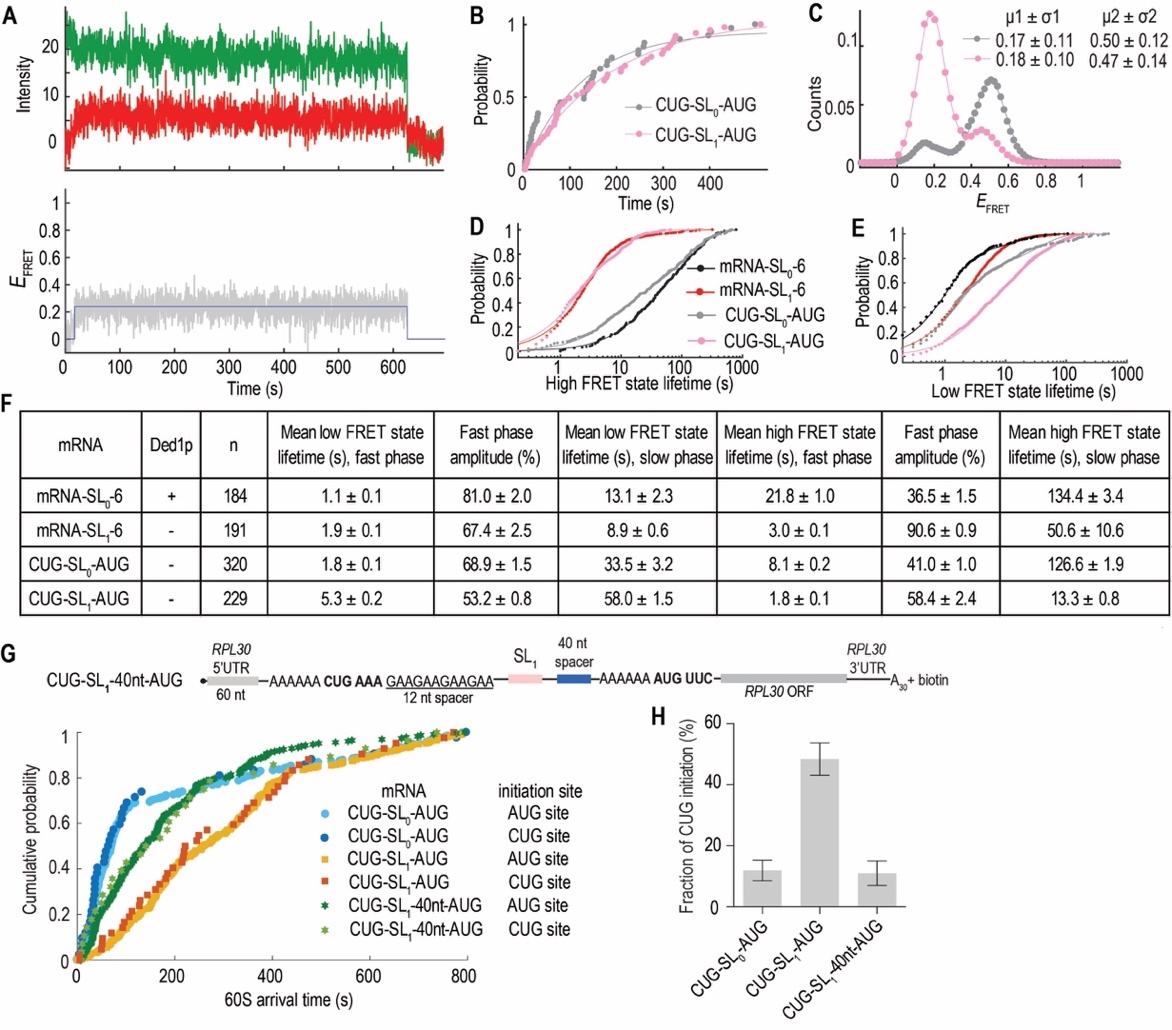

**Fig. S12. Stable hairpins in the 5′UTR stimulate upstream CUG initiation.**

(**A**) Sample trace of two-color mRNA–uS5 FRET that displays only a low mRNA–uS5 FRET state. (**B**) The cumulative probability distributions of the low FRET state lifetimes, from the traces showing only the low FRET state, observed for CUG-SL_0_-AUG (n = 39 out of 320, ~12% of the total traces) or CUG-SL_1_-AUG (n = 49 out of 229, ~21% of the total traces) mRNAs. Data were fit to a single-exponential equation to estimate the mean lifetimes (124 ± 25 s for CUG-SL_0_-AUG, 174 ± 14 s for CUG-SL_1_-AUG, mean ± 95% CI). (**C**) *E*_FRET_ distribution histograms from the mRNA–uS5 FRET experiments with CUG-SL_0_-AUG (n = 320) or CUG-SL_1_-AUG (n = 229) mRNAs. The distributions were fit to a double-Gaussian equation to obtain the mean *E*_FRET_ (μ) ± standard deviation (σ) values. The color scheme was the same as in (**B**). (**D**-**F**) Kinetic analysis of the mRNA–uS5 FRET fluctuations observed for mRNAs with or without a CUG codon upstream of the SL_0_ and SL_1_ sequences. The cumulative probability distributions of the observed high FRET state lifetimes (**D**) or low FRET state lifetimes (**E**) were fit to a double-exponential equation to estimate the mean lifetimes. Data in (**F**) were mean ± 95% CI. (**G**) Top, a cartoon illustration of the CUG-SL_1_-40nt-AUG mRNA, which contained a 40-nt unstructured spacer sequence between SL_1_ and the AUG codon. Bottom, the cumulative probability distributions of the observed 60S arrival time at AUG or CUG sites with varying mRNAs, measured in assay setup as in **Fig. 5B**. (**H**) Fraction of CUG initiation over total initiation events at both CUG and AUG sites. Data (mean ± 95% CI) for the CUG-SL_0_-AUG and CUG-SL_1_-AUG mRNAs are also shown in **Fig. 5E**, and for CUG-SL_1_-40nt-AUG mRNA, n= 254.

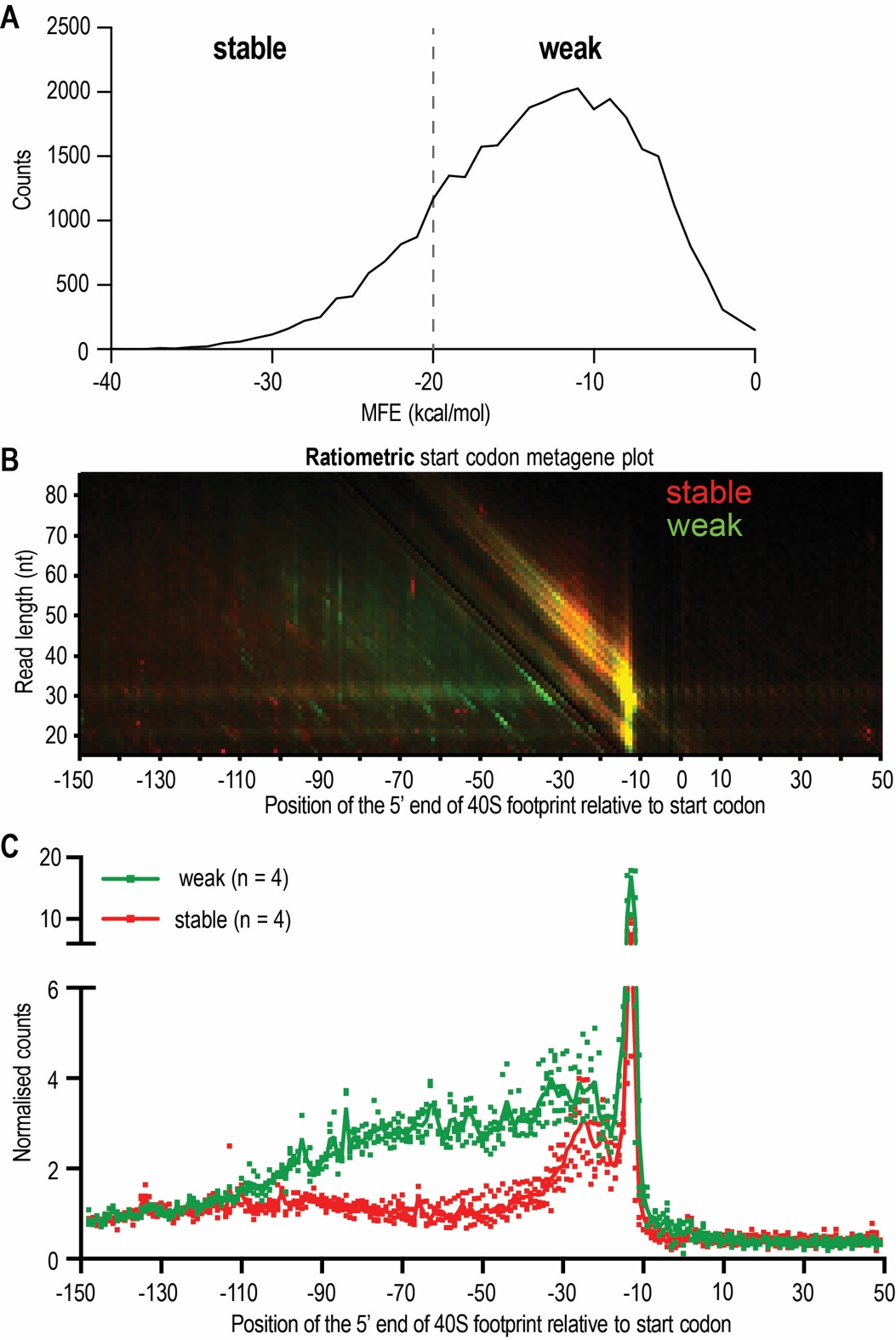

**Fig. S13. The 40S footprints in the 5′UTR adjacent to the start codons were suppressed by the presence of sequences with high propensity of secondary structure formation in human cells.** (**A**) The distribution of human mRNAs by the minimum free energy (MFE) for secondary structure formation predicted for a window between positions -55 to -5 relative to the start codons. Dashed line denotes the separation into the stable and weak structure buckets (see **Methods**). (**B**) Ratiometric start codon metagene plot of 40S footprints around the start codons in mRNAs with predicted weak (green) and stable (red) structures at positions -55 to -5. (**C**) Line graph of (**B**) showing data from four biological replicates.

**Table S1. Sequences of mRNAs and DNA oligonucleotides used in this study.**

| **mRNAs** | sequences (5' to 3', AUG start codon in green and UAA stop codon in red) |
| --- | --- |
| *RPL30* | m^7^GpppGGAAACAGACCGGAGUGUUUAAGAACCUACAGCUUAUUCAAUUAAUCAAUAUACGCAGAG**AUG**UUCCCAGUUAAAUCCCAAGAAUCUAUCAACCAAAAGUUGGCUUUGGUUAUCAAGUCUGGUAAGUACACCUUAGGUUACAAGUCCACUGUCAAGUCUUUGAGACAAGGUAAGUCUAAGUUGAUCAUCAUUGCCGCUAACACUCCAGUUUUGAGAAAGUCCGAAUUGGAAUAUUACGCUAUGUUGUCCAAGACUAAGGUCUACUACUUCCAAGGUGGUAACAACGAAUUGGGUACUGCUGUCGGUAAGUUAUUCAGAGUCGGUGUUGUCUCUAUUUUGGAAGCUGGUGACUCUGAUAUCUUGACCACCUUGGCU**UAA**AUAAGGUAAGUUCAAACGAUUUGUUGGAAGACAAUUGGUUUGAUGUGUAUUUUUCUAUAAUAUAUAAAUCUUAGACUAUUCAUUAACCUUAUUUUAUUUAAAAAAAAAAAAAAAAAAAAAAAAAAAAAA-biotin |
| *3xRPL30* | m^7^GpppGGAAACAGACCGGAGUGUUUAAGAACCUACAGCUUAUUCAAUUAAUCAAUAUACGCAGAGAAACAGACCGGAGUGUUUAAGAACCUACAGCUUAUUCAAUUAAUCAAUAUACGCAGAGAAACAGACCGGAGUGUUUAAGAACCUACAGCUUAUUCAAUUAAUCAAUAUACGCAGAG**AUG**UUCCCAGUUAAAUCCCAAGAAUCUAUCAACCAAAAGUUGGCUUUGGUUAUCAAGUCUGGUAAGUACACCUUAGGUUACAAGUCCACUGUCAAGUCUUUGAGACAAGGUAAGUCUAAGUUGAUCAUCAUUGCCGCUAACACUCCAGUUUUGAGAAAGUCCGAAUUGGAAUAUUACGCUAUGUUGUCCAAGACUAAGGUCUACUACUUCCAAGGUGGUAACAACGAAUUGGGUACUGCUGUCGGUAAGUUAUUCAGAGUCGGUGUUGUCUCUAUUUUGGAAGCUGGUGACUCUGAUAUCUUGACCACCUUGGCU**UAA**AUAAGGUAAGUUCAAACGAUUUGUUGGAAGACAAUUGGUUUGAUGUGUAUUUUUCUAUAAUAUAUAAAUCUUAGACUAUUCAUUAACCUUAUUUUAUUUAAAAAAAAAAAAAAAAAAAAAAAAAAAAAA+biotin |
| *SFT2* | m^7^GpppGGUAGUUUAGAGUAGGACUUAGAUUACCUGUAUUGUCUGCAGUUGCGUUUUUUUUUUUUGCUGGUAAAAAAAAAGAACAAGUGUGGAGAAU**AUG**UUCCCAGUUAAAUCCCAAGAAUCUAUCAACCAAAAGUUGGCUUUGGUUAUCAAGUCUGGUAAGUACACCUUAGGUUACAAGUCCACUGUCAAGUCUUUGAGACAAGGUAAGUCUAAGUUGAUCAUCAUUGCCGCUAACACUCCAGUUUUGAGAAAGUCCGAAUUGGAAUAUUACGCUAUGUUGUCCAAGACUAAGGUCUACUACUUCCAAGGUGGUAACAACGAAUUGGGUACUGCUGUCGGUAAGUUAUUCAGAGUCGGUGUUGUCUCUAUUUUGGAAGCUGGUGACUCUGAUAUCUUGACCACCUUGGCU**UAA**AUAAGGUAAGUUCAAACGAUUUGUUGGAAGACAAUUGGUUUGAUGUGUAUUUUUCUAUAAUAUAUAAAUCUUAGACUAUUCAUUAACCUUAUUUUAUUUAAAAAAAAAAAAAAAAAAAAAAAAAAAAAA+biotin |
| *PMA1* | m^7^GpppGGACCAAUAGUGAAAAUCUUUUUUUCUUCUAUAUCUACAAAAACUUUUUUUUUCUAUCAACCUCGUUGAUAAAUUUUUUCUUUAACAAUCGUUAAUAAUUAAUUAAUUGGAAAAUAACCAUUUUUUCUCUCUUUUAUACACACAUUCAAAAGAAAGAAAAAAAAUAUACCCCAGCUAGUUAAAGAAAAUCAUUGAAAAGAAUAAGAAGAUAAGAAAGAUUUAAUUAUCAAACAAUAUCAAU**AUG**UUCCCAGUUAAAUCCCAAGAAUCUAUCAACCAAAAGUUGGCUUUGGUUAUCAAGUCUGGUAAGUACACCUUAGGUUACAAGUCCACUGUCAAGUCUUUGAGACAAGGUAAGUCUAAGUUGAUCAUCAUUGCCGCUAACACUCCAGUUUUGAGAAAGUCCGAAUUGGAAUAUUACGCUAUGUUGUCCAAGACUAAGGUCUACUACUUCCAAGGUGGUAACAACGAAUUGGGUACUGCUGUCGGUAAGUUAUUCAGAGUCGGUGUUGUCUCUAUUUUGGAAGCUGGUGACUCUGAUAUCUUGACCACCUUGGCU**UAA**AUAAGGUAAGUUCAAACGAUUUGUUGGAAGACAAUUGGUUUGAUGUGUAUUUUUCUAUAAUAUAUAAAUCUUAGACUAUUCAUUAACCUUAUUUUAUUUAAAAAAAAAAAAAAAAAAAAAAAAAAAAAA+biotin |
| mRNA-SL_0_-6 | m^7^GpppGGAAACAGACCGGAGUGUUUAAGAACCUACAGCUUAUUCAAUUAAUCAAUAUACGCAGAGGAAGAAGAAGAAGAAGAAGAAGAAGAAGAAGAAGAAGAAGAAGAAGAAAAAA**AUG**UUCCCAGUUAAAUCCCAAGAAUCUAUCAACCAAAAGUUGGCUUUGGUUAUCAAGUCUGGUAAGUACACCUUAGGUUACAAGUCCACUGUCAAGUCUUUGAGACAAGGUAAGUCUAAGUUGAUCAUCAUUGCCGCUAACACUCCAGUUUUGAGAAAGUCCGAAUUGGAAUAUUACGCUAUGUUGUCCAAGACUAAGGUCUACUACUUCCAAGGUGGUAACAACGAAUUGGGUACUGCUGUCGGUAAGUUAUUCAGAGUCGGUGUUGUCUCUAUUUUGGAAGCUGGUGACUCUGAUAUCUUGACCACCUUGGCU**UAA**AUAAGGUAAGUUCAAACGAUUUGUUGGAAGACAAUUGGUUUGAUGUGUAUUUUUCUAUAAUAUAUAAAUCUUAGACUAUUCAUUAACCUUAUUUUAUUUAAAAAAAAAAAAAAAAAAAAAAAAAAAAAA+biotin |
| mRNA-SL_5_-6 | m^7^GpppGGAAACAGACCGGAGUGUUUAAGAACCUACAGCUUAUUCAAUUAAUCAAUAUACGCAGAGAGAAGUUACUUACUUCUAAAAAA**AUG**UUCCCAGUUAAAUCCCAAGAAUCUAUCAACCAAAAGUUGGCUUUGGUUAUCAAGUCUGGUAAGUACACCUUAGGUUACAAGUCCACUGUCAAGUCUUUGAGACAAGGUAAGUCUAAGUUGAUCAUCAUUGCCGCUAACACUCCAGUUUUGAGAAAGUCCGAAUUGGAAUAUUACGCUAUGUUGUCCAAGACUAAGGUCUACUACUUCCAAGGUGGUAACAACGAAUUGGGUACUGCUGUCGGUAAGUUAUUCAGAGUCGGUGUUGUCUCUAUUUUGGAAGCUGGUGACUCUGAUAUCUUGACCACCUUGGCU**UAA**AUAAGGUAAGUUCAAACGAUUUGUUGGAAGACAAUUGGUUUGAUGUGUAUUUUUCUAUAAUAUAUAAAUCUUAGACUAUUCAUUAACCUUAUUUUAUUUAAAAAAAAAAAAAAAAAAAAAAAAAAAAAA+biotin |
| mRNA-SL_4_-6 | m^7^GpppGGAAACAGACCGGAGUGUUUAAGAACCUACAGCUUAUUCAAUUAAUCAAUAUACGCAGAGAGUAAGCUACUUGCUUACUAAAAAA**AUG**UUCCCAGUUAAAUCCCAAGAAUCUAUCAACCAAAAGUUGGCUUUGGUUAUCAAGUCUGGUAAGUACACCUUAGGUUACAAGUCCACUGUCAAGUCUUUGAGACAAGGUAAGUCUAAGUUGAUCAUCAUUGCCGCUAACACUCCAGUUUUGAGAAAGUCCGAAUUGGAAUAUUACGCUAUGUUGUCCAAGACUAAGGUCUACUACUUCCAAGGUGGUAACAACGAAUUGGGUACUGCUGUCGGUAAGUUAUUCAGAGUCGGUGUUGUCUCUAUUUUGGAAGCUGGUGACUCUGAUAUCUUGACCACCUUGGCU**UAA**AUAAGGUAAGUUCAAACGAUUUGUUGGAAGACAAUUGGUUUGAUGUGUAUUUUUCUAUAAUAUAUAAAUCUUAGACUAUUCAUUAACCUUAUUUUAUUUAAAAAAAAAAAAAAAAAAAAAAAAAAAAAA+biotin |
| mRNA-SL_3_-6 | m^7^GpppGGAAACAGACCGGAGUGUUUAAGAACCUACAGCUUAUUCAAUUAAUCAAUAUACGCAGAGGUUAAGCCAUCUUGGCUUAACAAAAAA**AUG**UUCCCAGUUAAAUCCCAAGAAUCUAUCAACCAAAAGUUGGCUUUGGUUAUCAAGUCUGGUAAGUACACCUUAGGUUACAAGUCCACUGUCAAGUCUUUGAGACAAGGUAAGUCUAAGUUGAUCAUCAUUGCCGCUAACACUCCAGUUUUGAGAAAGUCCGAAUUGGAAUAUUACGCUAUGUUGUCCAAGACUAAGGUCUACUACUUCCAAGGUGGUAACAACGAAUUGGGUACUGCUGUCGGUAAGUUAUUCAGAGUCGGUGUUGUCUCUAUUUUGGAAGCUGGUGACUCUGAUAUCUUGACCACCUUGGCU**UAA**AUAAGGUAAGUUCAAACGAUUUGUUGGAAGACAAUUGGUUUGAUGUGUAUUUUUCUAUAAUAUAUAAAUCUUAGACUAUUCAUUAACCUUAUUUUAUUUAAAAAAAAAAAAAAAAAAAAAAAAAAAAAA+biotin |
| mRNA-SL_2_-6 | m^7^GpppGGAAACAGACCGGAGUGUUUAAGAACCUACAGCUUAUUCAAUUAAUCAAUAUACGCAGAGCCGAGCGGACCUCCUCGGCCGGAGACGGCCGAGGAGCAGGCGAGCCAAAAAA**AUG**UUCCCAGUUAAAUCCCAAGAAUCUAUCAACCAAAAGUUGGCUUUGGUUAUCAAGUCUGGUAAGUACACCUUAGGUUACAAGUCCACUGUCAAGUCUUUGAGACAAGGUAAGUCUAAGUUGAUCAUCAUUGCCGCUAACACUCCAGUUUUGAGAAAGUCCGAAUUGGAAUAUUACGCUAUGUUGUCCAAGACUAAGGUCUACUACUUCCAAGGUGGUAACAACGAAUUGGGUACUGCUGUCGGUAAGUUAUUCAGAGUCGGUGUUGUCUCUAUUUUGGAAGCUGGUGACUCUGAUAUCUUGACCACCUUGGCU**UAA**AUAAGGUAAGUUCAAACGAUUUGUUGGAAGACAAUUGGUUUGAUGUGUAUUUUUCUAUAAUAUAUAAAUCUUAGACUAUUCAUUAACCUUAUUUUAUUUAAAAAAAAAAAAAAAAAAAAAAAAAAAAAA+biotin |
| mRNA-SL_1_-6 | m^7^GpppGGAAACAGACCGGAGUGUUUAAGAACCUACAGCUUAUUCAAUUAAUCAAUAUACGCAGAGCCGAGCGGUCCUCCUCGGCCGGAGACGGCCGAGGAGGACCGCUCGGAAAAAA**AUG**UUCCCAGUUAAAUCCCAAGAAUCUAUCAACCAAAAGUUGGCUUUGGUUAUCAAGUCUGGUAAGUACACCUUAGGUUACAAGUCCACUGUCAAGUCUUUGAGACAAGGUAAGUCUAAGUUGAUCAUCAUUGCCGCUAACACUCCAGUUUUGAGAAAGUCCGAAUUGGAAUAUUACGCUAUGUUGUCCAAGACUAAGGUCUACUACUUCCAAGGUGGUAACAACGAAUUGGGUACUGCUGUCGGUAAGUUAUUCAGAGUCGGUGUUGUCUCUAUUUUGGAAGCUGGUGACUCUGAUAUCUUGACCACCUUGGCU**UAA**AUAAGGUAAGUUCAAACGAUUUGUUGGAAGACAAUUGGUUUGAUGUGUAUUUUUCUAUAAUAUAUAAAUCUUAGACUAUUCAUUAACCUUAUUUUAUUUAAAAAAAAAAAAAAAAAAAAAAAAAAAAAA+biotin |
| mRNA-SL_1_-46 | m^7^GpppGGAAACAGACCGGAGUGUUUAAGAACCUACAGCUUAUUCAAUUAAUCAAUAUACGCAGAGCCGAGCGGUCCUCCUCGGCCGGAGACGGCCGAGGAGGACCGCUCGGGAAGAAGAAGAAGAAGAAGAAGAAGAAGAAGAAGAAGAAGAAAAAA**AUG**UUCCCAGUUAAAUCCCAAGAAUCUAUCAACCAAAAGUUGGCUUUGGUUAUCAAGUCUGGUAAGUACACCUUAGGUUACAAGUCCACUGUCAAGUCUUUGAGACAAGGUAAGUCUAAGUUGAUCAUCAUUGCCGCUAACACUCCAGUUUUGAGAAAGUCCGAAUUGGAAUAUUACGCUAUGUUGUCCAAGACUAAGGUCUACUACUUCCAAGGUGGUAACAACGAAUUGGGUACUGCUGUCGGUAAGUUAUUCAGAGUCGGUGUUGUCUCUAUUUUGGAAGCUGGUGACUCUGAUAUCUUGACCACCUUGGCU**UAA**AUAAGGUAAGUUCAAACGAUUUGUUGGAAGACAAUUGGUUUGAUGUGUAUUUUUCUAUAAUAUAUAAAUCUUAGACUAUUCAUUAACCUUAUUUUAUUUAAAAAAAAAAAAAAAAAAAAAAAAAAAAAA+biotin |
| mRNA-SL_1_+20 | m^7^GpppGGAAACAGACCGGAGUGUUUAAGAACCUACAGCUUAUUCAAUUAAUCAAUAUACGCAGAG**AUG**UUCCCAGUUAAAUCCCAACCGAGCGGUCCUCCUCGGCCGGAGACGGCCGAGGAGGACCGCUCGGAAAAAAGAAUCUAUCAACCAAAAGUUGGCUUUGGUUAUCAAGUCUGGUAAGUACACCUUAGGUUACAAGUCCACUGUCAAGUCUUUGAGACAAGGUAAGUCUAAGUUGAUCAUCAUUGCCGCUAACACUCCAGUUUUGAGAAAGUCCGAAUUGGAAUAUUACGCUAUGUUGUCCAAGACUAAGGUCUACUACUUCCAAGGUGGUAACAACGAAUUGGGUACUGCUGUCGGUAAGUUAUUCAGAGUCGGUGUUGUCUCUAUUUUGGAAGCUGGUGACUCUGAUAUCUUGACCACCUUGGCU**UAA**AUAAGGUAAGUUCAAACGAUUUGUUGGAAGACAAUUGGUUUGAUGUGUAUUUUUCUAUAAUAUAUAAAUCUUAGACUAUUCAUUAACCUUAUUUUAUUUAAAAAAAAAAAAAAAAAAAAAAAAAAAAAA+biotin |
| CUGAAA-SL_0_-AUG | m^7^GpppGGAAACAGACCGGAGAGUUUAAGAACCUACAGCUUAUUCAAUUAAUCAAUAUACGCAGAGAAAAAA**CUG**AAAGAAGAAGAAGAAGAAGAAGAAGAAGAAGAAGAAGAAGAAGAAGAAGAAGAAGAAGAAGAAAAAA**AUG**UUCCCAGUUAAAUCCCAAGAAUCUAUCAACCAAAAGUUGGCUUUGGUUAUCAAGUCUGGUAAGUACACCUUAGGUUACAAGUCCACUGUCAAGUCUUUGAGACAAGGUAAGUCUAAGUUGAUCAUCAUUGCCGCUAACACUCCAGUUUUGAGAAAGUCCGAAUUGGAAUAUUACGCUAUGUUGUCCAAGACUAAGGUCUACUACUUCCAAGGUGGUAACAACGAAUUGGGUACUGCUGUCGGUAAGUUAUUCAGAGUCGGUGUUGUCUCUAUUUUGGAAGCUGGUGACUCUGAUAUCUUGACCACCUUGGCU**UAA**AUAAGGUAAGUUCAAACGAUUUGUUGGAAGACAAUUGGUUUGAUGUGUAUUUUUCUAUAAUAUAUAAAUCUUAGACUAUUCAUUAACCUUAUUUUAUUUAAAAAAAAAAAAAAAAAAAAAAAAAAAAAA+biotin |
| CUGAAA-SL_1_-AUG | m^7^GpppGGAAACAGACCGGAGAGUUUAAGAACCUACAGCUUAUUCAAUUAAUCAAUAUACGCAGAGAAAAAA**CUG**AAAGAAGAAGAAGAACCGAGCGGUCCUCCUCGGCCGGAGACGGCCGAGGAGGACCGCUCGGAAAAAA**AUG**UUCCCAGUUAAAUCCCAAGAAUCUAUCAACCAAAAGUUGGCUUUGGUUAUCAAGUCUGGUAAGUACACCUUAGGUUACAAGUCCACUGUCAAGUCUUUGAGACAAGGUAAGUCUAAGUUGAUCAUCAUUGCCGCUAACACUCCAGUUUUGAGAAAGUCCGAAUUGGAAUAUUACGCUAUGUUGUCCAAGACUAAGGUCUACUACUUCCAAGGUGGUAACAACGAAUUGGGUACUGCUGUCGGUAAGUUAUUCAGAGUCGGUGUUGUCUCUAUUUUGGAAGCUGGUGACUCUGAUAUCUUGACCACCUUGGCU**UAA**AUAAGGUAAGUUCAAACGAUUUGUUGGAAGACAAUUGGUUUGAUGUGUAUUUUUCUAUAAUAUAUAAAUCUUAGACUAUUCAUUAACCUUAUUUUAUUUAAAAAAAAAAAAAAAAAAAAAAAAAAAAAA+biotin |
| CUGAAA-SL_1_-40-AUG | m^7^GpppGGAAACAGACCGGAGAGUUUAAGAACCUACAGCUUAUUCAAUUAAUCAAUAUACGCAGAGAAAAAA**CUG**AAAGAAGAAGAAGAACCGAGCGGUCCUCCUCGGCCGGAGACGGCCGAGGAGGACCGCUCGGGAAGAAGAAGAAGAAGAAGAAGAAGAAGAAGAAGAAGAAGAAAAAA**AUG**UUCCCAGUUAAAUCCCAAGAAUCUAUCAACCAAAAGUUGGCUUUGGUUAUCAAGUCUGGUAAGUACACCUUAGGUUACAAGUCCACUGUCAAGUCUUUGAGACAAGGUAAGUCUAAGUUGAUCAUCAUUGCCGCUAACACUCCAGUUUUGAGAAAGUCCGAAUUGGAAUAUUACGCUAUGUUGUCCAAGACUAAGGUCUACUACUUCCAAGGUGGUAACAACGAAUUGGGUACUGCUGUCGGUAAGUUAUUCAGAGUCGGUGUUGUCUCUAUUUUGGAAGCUGGUGACUCUGAUAUCUUGACCACCUUGGCU**UAA**AUAAGGUAAGUUCAAACGAUUUGUUGGAAGACAAUUGGUUUGAUGUGUAUUUUUCUAUAAUAUAUAAAUCUUAGACUAUUCAUUAACCUUAUUUUAUUUAAAAAAAAAAAAAAAAAAAAAAAAAAAAAA+biotin |
| **DNA oligos for**  **mRNA hybridization** | sequences (5' to 3') |
| *RPL30* +20 Cy3-oligo | +CTTTTGG+TTGATA+GATT+C/3TYE563/ (+ indicates LNA nucleotide) |
| *RPL30* +31 Cy3-oligo | GATAACCAAAGCCAACTTTTGG-Cy3 |
| *RPL30* +45 Cy3-oligo | CTTACCAGACTTGATAACCAA-Cy3 |
| *RPL30* +55 Cy3-oligo | CTAAGGTGTACTTACCAGAC-Cy3 |
| *RPL30* +70 Cy3-oligo | CAGTGGACTTGTAACCTAAG-Cy3 |
| *PMA1* Cy3-oligo 1 | +CTTTTGG+TTGATA+GATT+C/3TYE563/ (+ indicates LNA nucleotide) |
| *PMA1* Cy3-oligo 2 | CTGGGAACATATTGATATTG-Cy3 |
| *PMA1* Cy3-oligo 3 | CAATGATTTTCTTTAACTAGCTGG-Cy3 |
| *PMA1* Cy3-oligo 4 | GATTGTTAAAGAAAAAATTTATCAACG-Cy3 |
| Kozak-RPL30-AUG blocking oligo | GGATTTAACTGGGAACATTTT |
| **Covalent labeling of mRNAs** | sequences (5' to 3') |
| fragment 2 for  splint ligation | /5Phos/rUrU rArArA rUrCrC rCrA/iCy3/ rArGrA rArUrC rUrArU rCrArA rCrCrA rArArA |
| fragment 3 for  splint ligation | 5'pGUUGGCUUUGGUUAUCAAGUCUGGUAAGUACACCUUAGGUUACAAGUCCACUGUCAAGUCUUUGAGACAAGGUAAGUCUAAGUUGAUCAUCAUUGCCGCUAACACUCCAGUUUUGAGAAAGUCCGAAUUGGAAUAUUACGCUAUGUUGUCCAAGACUAAGGUCUACUACUUCCAAGGUGGUAACAACGAAUUGGGUACUGCUGUCGGUAAGUUAUUCAGAGUCGGUGUUGUCUCUAUUUUGGAAGCUGGUGACUCUGAUAUCUUGACCACCUUGGCUUAAAUAAGGUAAGUUCAAACGAUUUGUUGGAAGACAAUUGGUUUGAUGUGUAUUUUUCUAUAAUAUAUAAAUCUUAGACUAUUCAUUAACCUUAUUUUAUUUAAAAAAAAAAAAAAAAAAAAAAAAAAAAAA+biotin |
| *RPL30*_fragment 1 | m^7^GpppGGAAACAGACCGGAGUGUUUAAGAACCUACAGCUUAUUCAAUUAAUCAAUAUACGCAGAGAUGUUCCCAG |
| *RPL30*_splint DNA | ACTTGATAACCAAAGCCAGATAGATTCTTGGGATTTACATCTCTGCGTATATTG |
| *3xRPL30*_fragment 1 | m^7^GpppGGAAACAGACCGGAGUGUUUAAGAACCUACAGCUUAUUCAAUUAAUCAAUAUACGCAGAGAAACAGACCGGAGUGUUUAAGAACCUACAGCUUAUUCAAUUAAUCAAUAUACGCAGAGAAACAGACCGGAGUGUUUAAGAACCUACAGCUUAUUCAAUUAAUCAAUAUACGCAGAGAUGUUCCCAG |
| *3xRPL30*_splint DNA | ACTTGATAACCAAAGCCAGATAGATTCTTGGGATTTACATCTCTGCGTATATTG |
| mRNA-SL_0_-6  _fragment 1 | m^7^GpppGGAAACAGACCGGAGUGUUUAAGAACCUACAGCUUAUUCAAUUAAUCAAUAUACGCAGAGGAAGAAGAAGAAGAAGAAGAAGAAGAAGAAGAAGAAGAAGAAGAAGAAAAAAAUGUUCCCAG |
| mRNA-SL_0_-6  _splint DNA | ACTTGATAACCAAAGCCAGATAGATTCTTGGGATTTACATTTTTTTCTTCTTCTTCTTC |
| mRNA-SL_2_-6  _fragment 1 | m^7^GpppGGAAACAGACCGGAGUGUUUAAGAACCUACAGCUUAUUCAAUUAAUCAAUAUACGCAGAGCCGAGCGGACCUCCUCGGCCGGAGACGGCCGAGGAGCAGGCGAGCCAAAAAAAUGUUCCCAG |
| mRNA-SL_2_-6  _splint DNA | ACTTGATAACCAAAGCCAGATAGATTCTTGGGATTTACATTTTTTTGGCTCGCCTGC |
| mRNA-SL_3_-6  _fragment 1 | m^7^GpppGGAAACAGACCGGAGUGUUUAAGAACCUACAGCUUAUUCAAUUAAUCAAUAUACGCAGAGGUUAAGCCAUCUUGGCUUAACAAAAAAAUGUUCCCAG |
| mRNA-SL_3_-6  _splint DNA | ACTTGATAACCAAAGCCAGATAGATTCTTGGGATTTACATTTTTTTGTTAAGCCAAG |
| mRNA-SL_4_-6  _fragment 1 | m^7^GpppGGAAACAGACCGGAGUGUUUAAGAACCUACAGCUUAUUCAAUUAAUCAAUAUACGCAGAGAGUAAGCUACUUGCUUACUAAAAAAAUGUUCCCAG |
| mRNA-SL_4_-6  _splint DNA | ACTTGATAACCAAAGCCAGATAGATTCTTGGGATTTACATTTTTTTAGTAAGCAAGTAG |
| mRNA-SL_5_-6  _fragment 1 | m^7^GpppGGAAACAGACCGGAGUGUUUAAGAACCUACAGCUUAUUCAAUUAAUCAAUAUACGCAGAGAGAAGUUACUUACUUCUAAAAAAAUGUUCCCAG |
| mRNA-SL_5_-6  _splint DNA | ACTTGATAACCAAAGCCAGATAGATTCTTGGGATTTACATTTTTTTAGAAGTAAGTAAC |

**References**

47.	S. F. Mitchell, S. E. Walker, M. A. Algire, E. H. Park, A. G. Hinnebusch, J. R. Lorsch, The 5′-7-methylguanosine cap on eukaryotic mRNAs serves both to stimulate canonical translation initiation and to block an alternative pathway. *Mol. Cell*. **39**, 950–962 (2010).

48.	E. H. Park, S. E. Walker, J. M. Lee, S. Rothenburg, J. R. Lorsch, A. G. Hinnebusch, Multiple elements in the eIF4G1 N-terminus promote assembly of eIF4G1•PABP mRNPs in vivo. *EMBO J.* **30**, 302–316 (2011).

49.	A. Petrov, R. Grosely, J. Chen, S. E. O’Leary, J. D. Puglisi, Multiple parallel pathways of translation initiation on the CrPV IRES. *Mol. Cell*. **62**, 92–103 (2016).

50.	O. Puig, B. Rutz, B. B. M. Luukkonen, S. Kandels-Lewis, E. Bragado-Nilsson, B. Séraphin, New constructs and strategies for efficient PCR‐based gene manipulations in yeast. *Yeast*. **14**, 1139–1146 (1998).

51.	Z. Zhou, P. Cironi, A. J. Lin, Y. Xu, S. Hrvatin, D. E. Golan, P. A. Silver, C. T. Walsh, J. Yin, Genetically encoded short peptide tags for orthogonal labelling by Sfp and AcpS phosphopantetheinyl transferases. *ACS Chem. Biol.* **2**, 337–346 (2007).

52.	M. R. Stark, J. A. Pleiss, M. Deras, S. A. Scaringe, S. D. Rader, An RNA ligase-mediated method for the efficient creation of large, synthetic RNAs. *RNA*. **12**, 2014–2019 (2006).

53.	J. W. van de Meent, J. E. Bronson, C. H. Wiggins, R. L. Gonzalez, Empirical Bayes Methods Enable Advanced Population-Level Analyses of Single-Molecule FRET Experiments. *Biophys. J.* **106**, 1327–1337 (2014).

54.	S. Peng, R. Sun, W. Wang, C. Chen, Single-molecule photoactivation FRET: a general and easy-to-implement approach to break the concentration barrier. *Angew. Chemie*. **129**, 6986–6989 (2017).

55.	E. F. Pettersen, T. D. Goddard, C. C. Huang, E. C. Meng, G. S. Couch, T. I. Croll, J. H. Morris, T. E. Ferrin, UCSF ChimeraX: Structure visualization for researchers, educators, and developers. *Protein Sci.* **30**, 70–82 (2021).

56.	R. Buschauer, Y. Matsuo, T. Sugiyama, Y. H. Chen, N. Alhusaini, T. Sweet, K. Ikeuchi, J. Cheng, Y. Matsuki, R. Nobuta, A. Gilmozzi, O. Berninghausen, P. Tesina, T. Becker, J. Coller, T. Inada, R. Beckmann, The Ccr4-Not complex monitors the translating ribosome for codon optimality. *Science*. **368**, eaay6912 (2020).

57.	A. R. Gruber, R. Lorenz, S. H. Bernhart, R. Neuböck, I. L. Hofacker, The Vienna RNA Websuite. *Nucleic Acids Res.* **36**, W70–W74 (2008).
